# Elimination of myotonia improves myopathy in a muscleblind knockout model of myotonic dystrophy

**DOI:** 10.1101/2025.09.19.677400

**Authors:** Matthew T. Sipple, Sakura A. Hamazaki, Vanessa Todorow, Lily A. Cisco, Katherine M. Lupia, Christina S. Heil, Peter Meinke, Charles A. Thornton, John D. Lueck

## Abstract

A cardinal sign of myotonic dystrophy type 1 (DM1) is slow of muscle relaxation after voluntary contraction known as myotonia. Myotonia results from mis-regulated splicing of chloride channel 1 (ClC-1), leading to loss of channel function and runs of involuntary action potentials in muscle fibers. Heralding the onset of weakness, myotonia is often the first symptom of DM1, and raising the possibility that muscle hyperexcitability promotes the subsequent development of myopathy. We used genome editing to test this possibility by deleting the alternatively spliced and frameshift inducing ClC-1 exon 7a (E7a) in the Mbnl1 knockout model of DM1. Although several ClC-1 exons exhibit mis-regulated splicing in DM1, deletion of this single cryptic exon was sufficient to restore ClC-1 function and eliminate myotonia systemically and permanently. As determined by long-read sequencing, deletion of E7a reduced the frequency of other splicing defects in ClC-1 transcripts, likely as a passive consequence of restoring reading frame and nonsense surveillance. Furthermore, we observed significantly improved muscle force generation, fiber-type distribution, and histology, and partial restoration of the muscle transcriptome, including differential gene expression and alternative splicing, in non-myotonic Mbnl1 knockout mice. These results suggest that E7a inclusion is a lynchpin splice event that contributes to skeletal myopathy, highlighting myotonia as a therapeutic target and an outcome of interest in DM1.

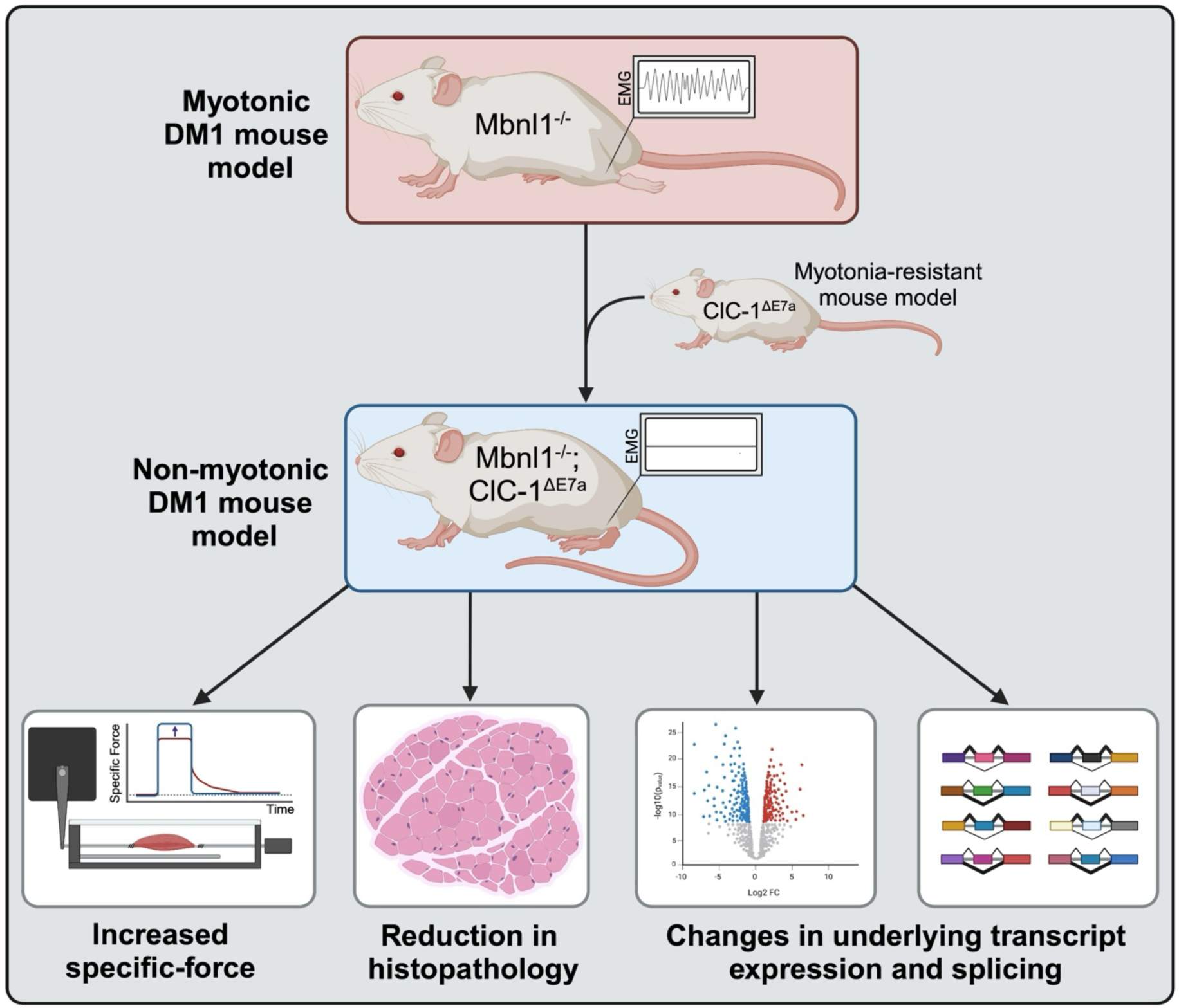

## Introduction

Myotonic dystrophy type 1 (DM1) is an autosomal dominant, multi-system disorder caused by trinucleotide repeat (CTG)_n_ expansion in the DM1 protein kinase (*DMPK*) gene^1^. Transcripts harboring expanded (CTG)_n_ repeats form nuclear foci that sequester mRNA binding proteins, such as the muscleblind-like (MBNL) family of splicing factors. Loss of active MBNL proteins, that normally coordinate many alternative splice events, results in widespread changes in transcript isoforms expressed and consequently progressive muscle degeneration and myotonia, the impaired relaxation of skeletal muscles following voluntary contraction.

Dysregulated splicing of *CLCN1*, that encodes the main voltage-sensitive chloride channel (ClC-1) in skeletal muscle, has been directly linked to myotonia^2,3^. Increased inclusion of the alternative exon 7a (E7a) in Clcn1 transcripts in DM1 causes a frameshift that leads to a premature termination codon (PTC), and thus a non-functional truncated ion channel as well as non-sense mediated decay (NMD) of the mRNA^4^. This process is conserved across both murine and human homologs. Loss of ClC-1 current, that normally dampens myofiber excitability during repetitive stimulation, produces prolonged runs of action potentials that follow voluntary muscle activation and clinically manifests as myotonia^5^.

Although DM1 represents a hypervariable disease, myotonia is a distinguishing characteristic that differentiates it from other muscular dystrophies. Myotonia often presents early in DM1 disease progression with a significant impact on muscle function that precedes the onset of progressive muscle weakness and degeneration^6^. Prior studies that have investigated the potential direct role of muscle hyperexcitability in exacerbating muscle weakness in DM1 have provided mixed results. Pre-clinical studies have identified the potential for myotonia to drive changes in fiber-type distributions to more oxidative types that output less maximal specific force, possibly contributing to weakness^7,8^. Comparative studies with models of myotonia congenita (mutations in *CLCN1*) have also shown the capacity to drive changes in transcript expression^9^. Moreover, our group has previously demonstrated that only myotonia and altered calcium channel splicing was sufficient to produce severe muscle weakness in murine models^10^. Despite these pre-clinical studies showing promise in myotonia as a driving factor in myopathy, a clinical trial with mexiletine, a repurposed anti-myotonic agent, found there to be no impact on clinical tests for muscle function, including the 6-minute walk time or grip strength after 6-months of treatment. This trial, however, was limited in the capacity to achieve sustained suppression of myotonia and relatively short duration of treatment^11^.

Accordingly, the status of myotonia as a driver of myopathy in DM1 and the value of myotonia suppression as a disease-modifying strategy requires further investigation. This is of particular importance as myotonia has emerged as a main clinical outcome for assessing the success of novel therapeutics in DM1, such as the phase III HARBOR trial investigating the effects of Delpacibart etedesiran—an antibody-oligonucleotide conjugate designed to knockdown *DMPK* expression (NCT06411288). Here, to definitively isolate the role of long-term exposure to myotonia in the development of DM1 myopathy, we developed a novel myotonia resistant mouse model through genomic deletion of *Clcn1* E7a (ClC-1^ΔE7a^); the exclusion of E7a in *Clcn1* transcripts with ASOs has been demonstrated to eliminate myotonia in DM1 mouse models^12^. By crossing the ClC-1^ΔE7a^ mouse model with the Mbnl1^−/−^ model of DM1, we eliminated myotonia systemically throughout its lifespan. This afforded the direct comparison of muscle physiology via *in vitro* and *in vivo* muscle contraction, histopathology (e.g., central nucleation), and the transcriptome between myotonic (Mbnl1^−/−^) and non-myotonic (Mbnl1^−/−^ ; ClC-1^ΔE7a^) mice. Overall, these mice feature muscular function closer to wildtype mice with significant reduction in aberrant transcript expression and splicing and DM1 muscle histopathology.

## Results

### Generation of a *Clcn1* exon 7a deletion mouse for elimination of myotonia from DM1 mouse models

To model the effects of the ideal, systemic long-term anti-myotonic treatment, we used CRISPR/Cas9 to excise E7a and flanking intronic regions from the *Clcn1* gene (Fig. 1a, Fig. S1a). ClC-1^ΔE7a/ΔE7a^ mice were thus incapable of E7a inclusion in transcripts regardless of DM1 mechanisms of aberrant splicing, which has previously been shown to be successful in completely eliminating myotonia with spice-shifting ASOs^8,12^. We chose to cross these mice with the Mbnl1^−/−^ model of DM1 in this study, as they feature a consistent and well characterized muscle phenotype, including robust myotonia, through global deletion of the main Mbnl isoform in skeletal muscle that is sequestered in DM1^13^.

**Figure 1.**
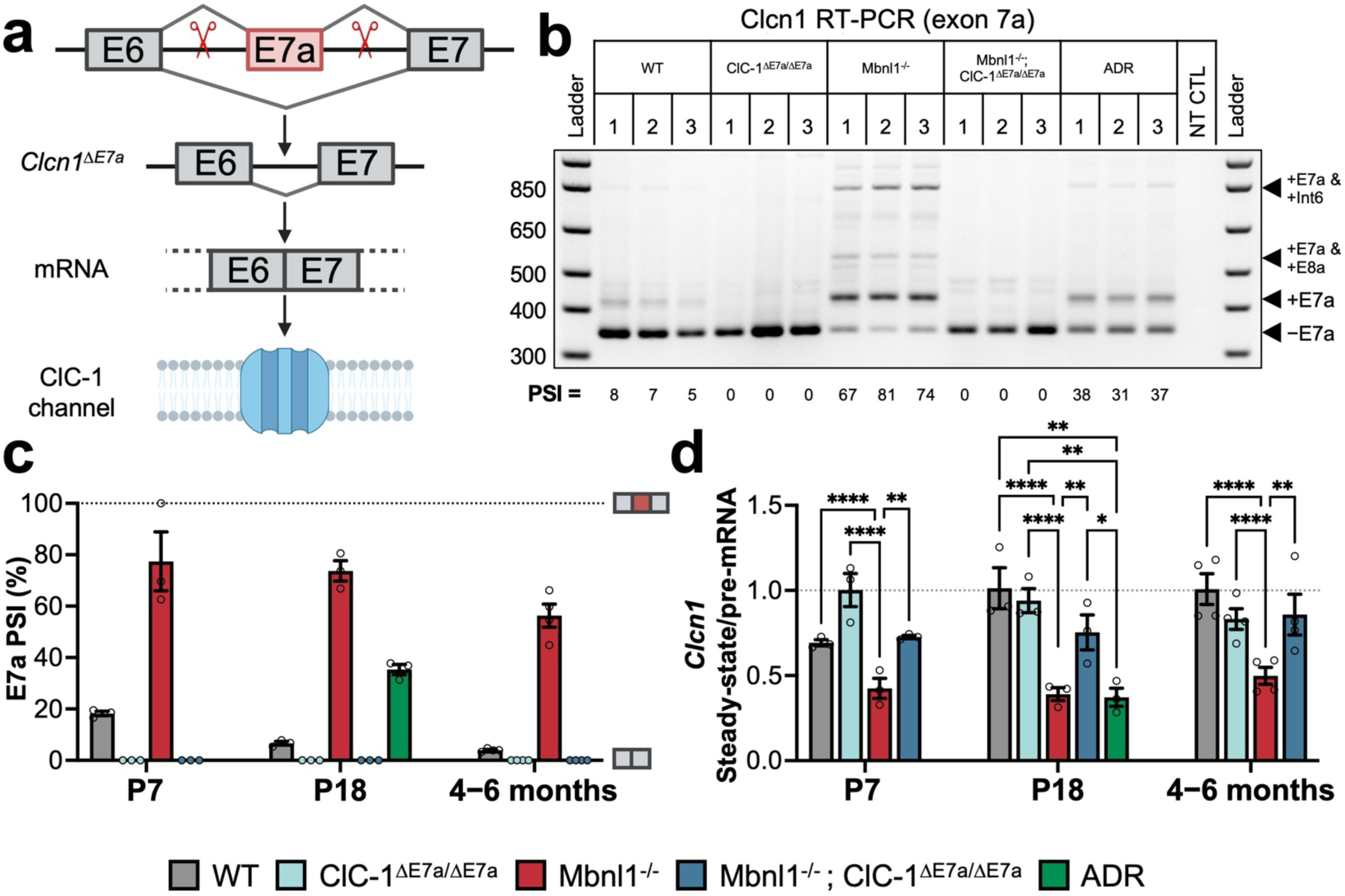
Generating a forced exon 7a exclusion mouse model that corrects changes *Clcn1* splicing and RNA stability in Mbnl1 knockout mice. (a) Schematic of CRISPR/Cas9 approach to develop a myotonia-resistant mouse model by excision of the exon 7a sequence from the *Clcn1* gene. Scissors represent approximate location of the gRNA sequences in the intronic regions flanking the exon 7a (sequences provided in supplemental figure 1). (b) Ethidium bromide-stained agarose gel of RT-PCR products generated with primers flanking exon 7a from RNA isolated from quadriceps muscle from WT, ClC-1^ΔE7a/ΔE7a,^ Mbnl1^−/−^, Mbnl1^−/−^ ; ClC-1^ΔE7a/ΔE7a,^ and ADR (NMD control) mice (P18). Each lane represents an individual mouse, with biological replicates denoted by number. Ladder (1kb Plus ladder, Invitrogen) and no template control (NT CTL) shown. Unique transcript isoforms identified by the distinct molecular weight of the amplicon indicated on the right. –E7a represents the adult, wildtype isoform. (c) Quantitation of exon 7a percent spliced in (PSI) based on relative densitometry of the amplicons shown in panel (b), as well as for samples at P7 and 4-6-months of age, with bars shown mean ± SE and individual biological replicates indicated by dots (n=3-4/genotype/timepoint, other agarose gels shown in Fig. S2a). (d) To determine relative RNA stability, the ratio of relative *Clcn1* steady-state mRNA to pre-mRNA was determined with the use of separate RT-qPCR reactions with primer-probe sets directed at constitutively spliced exon-exon boarders and intron-exon boarders, respectively. Steady-state and pre-mRNA experiments were each completed with the same reference *Tbp* primer-probe set and normalized to the average of P18 WT samples to allow for the determination of the ratio of relative *Clcn1* steady-state expression / *Clcn1* pre-mRNA (n = 3-4 biological replicates/genotype/timepoint; completed in technical triplicate). Dots represent biological replicates and bars plotted mean ± SE. Statistical comparisons were completed with two-way with Tukey’s multiple comparisons (only significant intra-timepoint comparisons shown for simplicity). Significant comparisons indicated as multiplicity-adjusted p-value: * p < 0.05, ** p < 0.01, *** p < 0.001, and **** p < 0.0001.

RT-PCR experiments were conducted on total RNA isolated from quadriceps muscle of WT, ClC-1^ΔE7a/ΔE7a^, Mbnl1^−/−^, and Mbnl1^−/−^ ; ClC-1^ΔE7a/ΔE7a^ mice, with primers annealing in exons flanking this alternatively spliced region to assess successful removal of E7a from *Clcn1* transcripts of ClC-1^ΔE7a/ΔE7a^ and Mbnl1^−/−^ ; ClC-1^ΔE7a/ΔE7a^ mice. To control for the effect of NMD on *Clcn1* transcripts, we used *adr*^mto2j^ (arrested development of righting response; ADR) mice, that have a single-nucleotide deletion in the *Clcn1* gene resulting in a total loss of ClC-1 ^4^. For ClC-1^ΔE7a/ΔE7a^ and Mbnl1^−/−^ ; ClC-1^ΔE7a/ΔE7a^ mice, no products with molecular weights consistent with E7a containing isoforms were observed, as shown by the representative gel of P18 muscle samples in Figure 1b. Mbnl1^−/−^ muscle had a significantly increased proportion of isoforms containing E7a (73.7 ± 3.97%) as compared to wildtype (6.65 ± 0.71%; expressed by percent-spliced-in, PSI) and quantified in figure 1c with additional early post-natal (P7) and aged (4-6-months-old) timepoints (Representative gels shown Fig. S2a).

E7a inclusion causes a shift in the downstream reading frame and subsequent NMD of E7a inclusion transcripts; this was shown to decrease steady-state concentrations of *Clcn1* mRNA^4^. We used RT-qPCR to assess recovery in *Clcn1* RNA stability (ratio of steady-state to pre-mRNA) in the setting of E7a excision with primer-probe sets designed to target exon-exon boarders and intron-exon boarders in *Clcn1*, to provide relative abundances of *Clcn1* mature steady-state mRNA and pre-mRNA, respectively (Fig. S1b, S1c). Decreased mature-to pre-mRNA ratios were observed for Mbnl1^−/−^ samples at all time points when compared to age matched WT mice (p<0.0001). Accordingly, the lower ratio of mature- to pre-mRNA indicates an overall reduction in *Clcn1* mRNA stability (e.g., increased rate of degradation through processes such as NMD of transcripts). Mbnl1^−/−^ ; ClC-1^ΔE7a/ΔE7a^ samples had ratios significantly greater than Mbnl1^−/−^ and equivalent to WT samples at P7, P18 and the 4-6-month timepoints (p-value of 0.0018, 0.0219, and 0.0018, respectively for Mbnl1^−/−^ vs. Mbnl1^−/−^ ; ClC-1^ΔE7a/ΔE7a^). Moreover, these changes in mRNA stability closely trend with the changes in E7a splicing (Fig. S2b). Additionally, *Clcn1* transcripts from ADR mice (P18) had a significantly decreased ratio of mature to pre-mRNA compared to wildtype controls (Fig. 1d), due to the frame-shift mutation resulting in NMD of all transcripts regardless of splice pattern.

### Long-read, whole-transcript sequencing revealed changes in *Clcn1* transcript isoform utilization in the setting of exon 7a deletion, Mbnl1 depletion, and constitutive NMD

Despite the longstanding use of conventional RT-PCR coupled with gel electrophoresis to assess changes in single splice events, or read RNA-seq that allows for the assessment of numerous isolated splice events, information is lost on the identity of the full transcript isoforms. This is particularly important for genes with numerous alternatively spliced regions, and thus a multitude of possible underlying transcript isoforms, such as for *Clcn1*. In DM1, in addition to E7a retention, increased retention of intron 2, intron 6, and inclusion of exon 8a, among others have also been observed, as illustrated in Figure 2a^2,3^. However, rather unexpectedly in the setting of these various other splicing changes seen in DM1, correction of only E7a splicing with ASOs was sufficient to eliminate myotonia and completely rescue ClC-1 channel function in DM1 mouse models^12^. Accordingly, this brings to question whether each of these splice events are independently regulated by Mbnl1 or a single event is controlled by Mbnl1, and other events are enriched as a byproduct of changes in the aggregate mRNA stability. Outside of *Clcn1*, prior studies have also suggested the role of cis-acting intronic features to direct other splicing events in transcripts, and thus E7a splicing, could have served as keystone splicing event that regulates the other splice events upregulated in DM1^14^.

**Figure 2.**
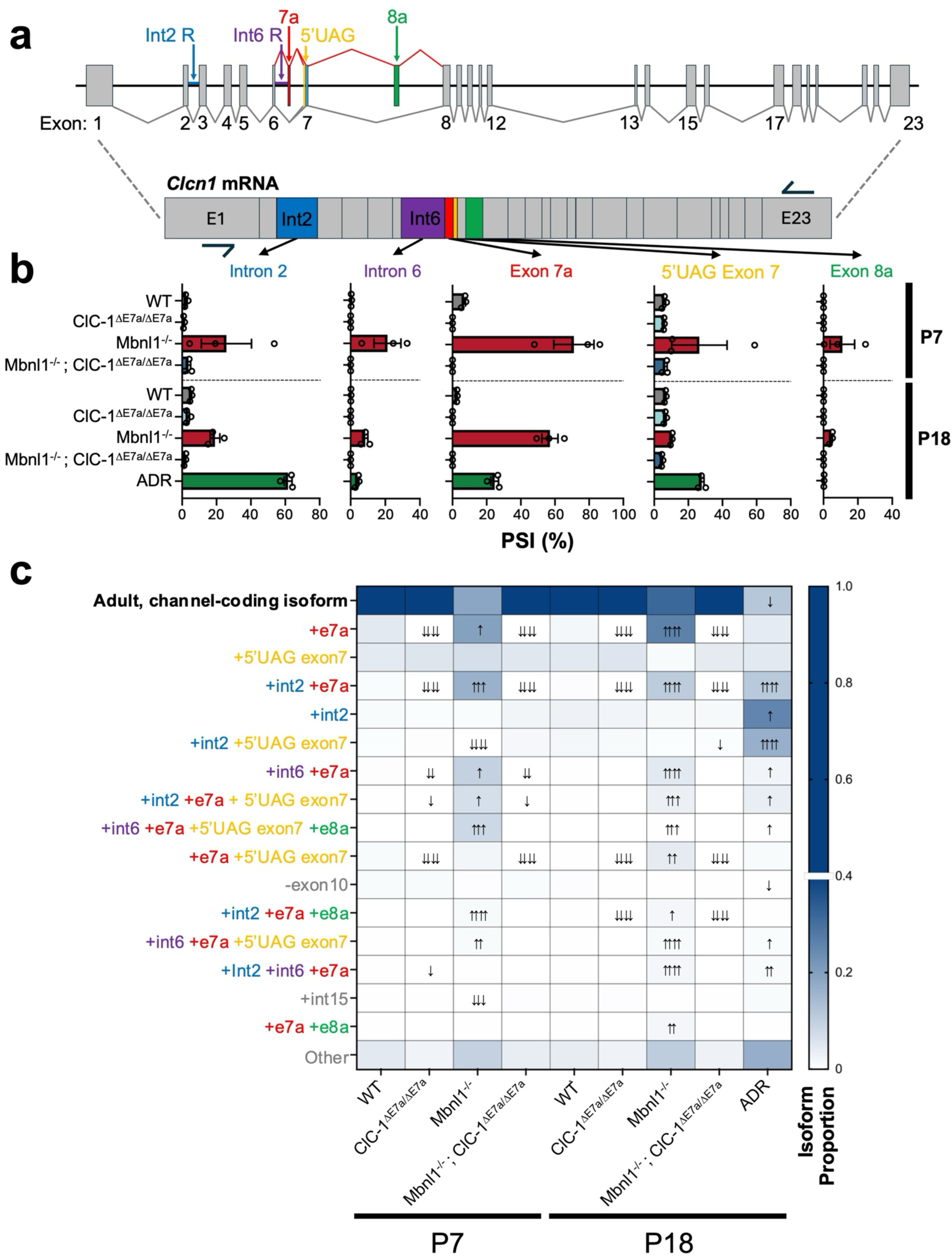
Whole-transcript sequencing illustrates Mbnl1 dependent and independent splicing events in the setting of NMD causing splicing changes. (a) Representation of the murine *Clcn1* gene with constitutive exons (gray) and common alternative splice elements seen in DM1 annotated in distinct colors. Downward projection represents a schematic of post-splicing *Clcn1* mRNA containing all the above annotated alternative splicing features not typically present in the adult, ion-channel coding isoform of the transcript. Approximate forward and reverse primer locations—exon 1 and exon 23, respectively-–used for generating the whole-*Clcn1* coding region amplicons for long-read sequencing libraries shown as arrows. Whole-*Clcn1* amplicon sequencing libraries were generated for both early (P7) and late (P18) post-natal mice with RNA isolated from quadriceps muscle of WT, ClC-1^ΔE7a/ΔE7a,^ Mbnl1^−/−^, and Mbnl1^−/−^ ; ClC-1^ΔE7a/ΔE7a^ genotypes as well as late post-natal ADR samples used as the NMD control (n = 3 mice/genotype/timepoint). To determine transcript isoform utilization for each sample, reads were aligned to *Ch. 6* (mm39) and collapsed into isoforms with greater than 100 supporting reads and the most abundant reads were annotated for quantification with the use of FLAIR. (b) Quantification of prominent individual splice elements, as expressed as percent spliced in (PSI), based on whole-length transcript reads. Bars shown mean ± SE with individual biological replicates shown as dots. (c) Assessment of transcript isoform utilization with FLAIR with reads quantified for the adult, protein-coding isoform, and the 15 most abundant alternate isoforms seen in the library, shown as a heat map; rows indicate transcript isoform based on the element or combination of elements that differentiate from the adult, protein-coding isoform and columns indicate the different genotype/timepoint conditions assessed. Color intensity of each box represents the average proportion of that isoform in the pool of reads for each sample (n = 3 mice/genotype/timepoint), as per the scale bar to right (note, the broken scale skewed to lower isoform proportions to visually highlight differences in the lower abundance alternative transcript isoforms). Arrows superimposed on boxes represent statistical comparisons between each genotype/timepoint and the age-matched wild-type samples using DESeq2; number of arrows indicates significance level (p < 0.05 ↑, p < 0.01 ↑↑, p < 0.001 ↑↑↑, or p < 0.0001 ↑↑↑↑) and direction indicates up/down regulated compared to age-matched wild-type samples.

To address this, we used long-read PacBio sequencing that provided contiguous reads of the entire possible protein coding region of *Clcn1* transcripts from RNA extracted from early and late postnatal mice (Fig. S3a). By analyzing full-length reads of the *Clcn1* transcripts we were able to observe changes in individual splice events within a transcript (Fig. 2b), as well as changes in unique transcript isoforms made up of different combinations of these splice events (Fig. 2c). Mbnl1^−/−^ mice at both P7 and P18 displayed a higher PSI for E7a as well as intron 2 and intron 6, as compared to age matched WT mice, and these values correlated with those determined by traditional RT-PCR (Fig. S3b, S3c). Interestingly, however, the NMD control samples (ADR) also altered PSIs of these events despite featuring no direct manipulations of their endogenous Mbnl1 proteins (Fig. 2b). Therefore, by inspecting only isolated splice events it is difficult to differentiate effects of shifts in splicing that are secondary to changes the overall stability of the mRNA pool or from direct alterations in Mbnl1.

To inspect changes in *Clcn1* isoform utilization, we used an established pipeline (FLAIR), that identifies and quantifies the use of different transcript isoforms based on contiguous long reads. A heatmap is shown with the columns representing the mean of each group (N=3) per genotype and timepoint and the rows representing unique *Clcn1* isoforms identified and denoted by features that differ from the canonical adult isoform (Fig. 2c). In both early (P7) and late postnatal (P18) Mbnl1^−/−^ samples, there was predominantly upregulation of transcript isoforms that include E7a, including isoforms that only feature E7a inclusion as well as combinations of E7a coupled with intron 2 inclusion, intron 6 inclusion, and/or exon 8a, all represent potential NMD candidates. However, isoforms with these other splicing events in isolation, such as isolated intron 2 retention (row 4), were not differentially expressed as compared to age matched controls. Interestingly, in the P18 ADR samples, which feature a shift in the reading frame for all transcripts, and thus NMD candidates regardless of splicing patterns exhibited an increase in abundance of transcripts containing various combinations of E7a inclusion, intron 2 inclusion, and intron 6 inclusion as well as others, despite unperturbed Mbnl1 function. Accordingly, upregulation of many of these splicing events is likely secondary to changes in the stability of the dominant isoforms in the pool rather than direct impacts of Mbnl1. This was further illustrated when the splicing events were assessed in a combinatorial manner by sorting the transcript pool for those with and without intron 2 inclusion or E7a inclusion; for Mbnl1^−/−^ E7a inclusion was greater in intron 2 containing isoforms as compared to transcripts lacking intron 2, whereas E7a retention was independent of intron 2 retention in ADR samples (Fig. S3d). The reciprocal relationship of intron 2 PSI based on E7a status also showed similar trends (Fig. S3e).

### Exon 7a deletion restored ClC-1 channel function and eliminated myotonia in Mbnl1 knockout mice

Patch-clamp electrophysiology represents the gold standard to assess functional protein levels of ion channels, such as ClC-1^15^. Therefore, to determine if genetic elimination of the E7a sequence from *Clcn1* was sufficient to recover wildtype levels of functional recovery of ClC-1, we collected whole-cell voltage-clamp recordings of ClC-1 currents from myofibers isolated from WT, ClC-1^ΔE7a/ΔE7a^, Mbnl1^−/−^, and Mbnl1^−/−^ ; ClC-1^ΔE7a/ΔE7a^ flexor digitorum brevis (FDB) muscles. Representative traces of current-densities collected from fibers at postnatal day 18 are shown in Figure 3a in response to standard volage-clamp protocol^16,17^. The traces display the characteristic deactivation pattern and inward rectification of ClC-1 channels^4,16–18^. Furthermore, through inspecting both the peak inward instantaneous current as well as the average steady-state current, we were able to assess the current-voltage (I-V) relationship in these different mice (Fig. 3b,c). Peak instantaneous inward current density at the onset of the hyperpolarization voltage step to −140 mV in Mbnl1^−/−^ myofibers had a significantly reduced current density (− 25.3±1.2 pA/pF; n = 16 fibers) compared to wildtype (−42.1 ± 1.7 pA/pF; n = 14 fibers; p < 0.0001). Whereas the current density of Mbnl1^−/−^ ; ClC-1^ΔE7a/ΔE7a^ mice was (−56.4 ± 2.7 pA/pF; n = 14 fibers) greater than wildtype, representing at minimum a complete rescue of ClC-1 function. ADR controls displayed negligible ClC-1 current densities (0.03 ± 0.06 pA/pF; n = 7 fibers), as expected (Fig. 3d). These findings were largely consistent with those seen at post-natal day 12 and voltage-dependence of activation was largely independent of genotype^4^. Moreover, ClC-1^ΔE7a/ΔE7a^ displayed current densities greater than WT across the entire early postnatal period assessed (P7-P18; Fig. S4). Linear properties and fitting parameters summarized in Supplemental Table 2.

**Figure 3.**
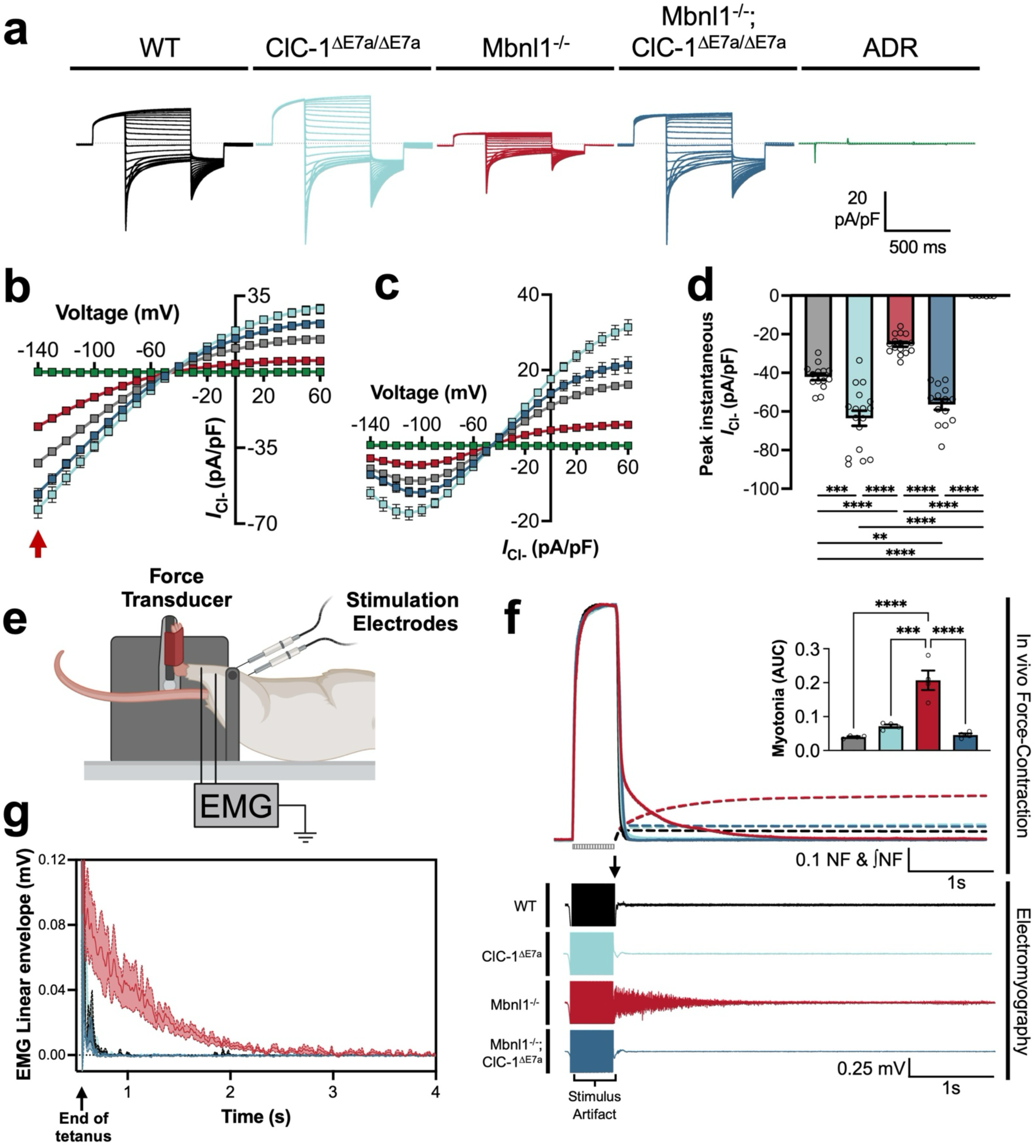
Forced exclusion of exon 7a in Mbnl1 knockout mice corrects ClC-1 channel function to wildtype levels and eliminates myotonia. (a) Whole-cell patch-clamp recordings of ClC-1 currents from FDB myofibers isolated from WT, ADR, ClC-1^ΔE7a/ΔE7a,^ Mbnl1^−/−^, and Mbnl1^−/−^ ; ClC-1^ΔE7a/ΔE7a^ mice (P18), representative traces shown (current expressed as current density, normalized to myofiber capacitance, scale bar shown). Recordings were collected with solutions designed to isolate ClC-1 currents with the use of the specific blocker 9-AC and offline subtraction to exclude leak and capacitive currents. Voltage protocol consisted of an initial depolarization to +60mV for maximal channel activation followed by a voltage step from −140 mV to +60 mV (Δ10mV), and then a step to −100 mV for tail-current analysis before returning to resting (−50 mV). Recordings were collected from multiple mice for each genotype yielding 7-17 recordings per group. (b) IV-plot of instantaneous current-density determined as the peak inward current at the start of the voltage step. Shown mean ± SE for each voltage-step. Connecting line represents fit to modified-Boltzmann equation. (c) IV-plot of steady-state current-density representing the average current density at the end of each voltage step. (d) Peak inward instantaneous current-density at −140 mV (red arrow in panel b). Bars shown mean ± SE with replicates shown as dots. (e) Schematic of experimental setup for collecting paired plantarflexion force recordings and electromyograms (EMGs) from anesthetized mice. Stimulating electrodes labeled positioned on medial aspect of the proximal ipsilateral lower limb for stimulation of the tibial nerve. EMG electrodes placed subcutaneously in proximal and distal aspects of the posterior compartment. (f) Representative normalized force traces from WT, ClC-1^ΔE7a/ΔE7a,^ Mbnl1^−/−^, and Mbnl1^−/−^ ; ClC-1^ΔE7a/ΔE7a^ mice are shown in response to a 500 ms (150 Hz) stimulus (represented by the box with vertical lines). To quantify the force generated after the end of the stimulus, integration of the force-time curve was completed, starting at the end of the tetanic stimulus (as shown by dashed lines). Total area under the curve was then compared for recordings from each of the groups (inset). Representative EMG recordings shown below the force traces and aligned with the tetanic stimulus above. Stimulus artifact as indicated on graph, with magnitude maxed at ± 0.5 mV to provide adequate scale for visualization of post-stimulatory electrical activity—i.e., myotonia. Scales as shown. (g) Averaged linear envelope of EMG myotonia generated by rectifying the raw signals and applying a second-order Butterworth low-pass filter (n = 4 mice/genotype). Arrow indicates the approximate alignment in time as indicated by the arrow on the representative traces in panel f, time starting at 0.5 ms (the end of the tetanic stimuli). Data shown as mean (solid line) ± SE as shown by (dotted lines with shading within 1 SE of the mean). Statistical comparisons shown in panels d and f, represent one-way ANOVA with Tukey’s multiple comparisons. Significance indicated as multiplicity adjusted p-value < 0.05 *, 0.01 **, 0.001 ***, 0.0001 ****.

From initial observation of Mbnl1^−/−^ ; ClC-1^ΔE7a/ΔE7a^ mice cage behavior, it was clear that our goal of eliminating myotonia from this DM1 mouse line was successful. No Mbnl1^−/−^ ; ClC-1^ΔE7a/ΔE7a^ mouse observed during of any of the experiments described here displayed any of the characteristic hindlimb stiffness when physical stimulation was applied (professional observation, not formally quantified), whereas all Mbnl1^−/−^ displayed characteristic hind limb stiffness that has been previously described^13^. Interestingly, Mbnl1^−/−^ ; ClC-1^ΔE7a/+^ mice with only a single allele with forced E7a exclusion, also lacked any evidence of myotonia, and thus for all physiologic experiments both heterozygote and homozygote ClC-1^ΔE7a^ mice were used as indicated by either ClC-1^ΔE7a^ or Mbnl1^−/−^ ; ClC-_1ΔE7a._

To provide a more quantitative assessment of myotonia, we utilized *in vivo* plantarflexion of anesthetized mice paired with electromyography (EMG); this allows for assessment of both mechanical myotonia (i.e. delayed muscle relaxation following voluntary contraction, akin to clinical myotonia) and electrical myotonia (i.e., continue firing of action potentials after stimulation through the neuromuscular junction (NMJ)). Force recordings were collected in response to a tetanic stimulus delivered through the tibial nerve, and then a subsequent tetanic stimulus was delivered with monopolar electrodes placed subcutaneously along the posterior compartment to record underlying electrical activity, as shown by the schematic in Figure 3e. Mechanical myotonia was quantified by integrating the area under the curve (AUC) of normalized force versus time trace immediately after the cessation of the stimulus. Mbnl1^−/−^ displayed the continued generation of force, as shown by the representative trace in Figure 3f, as well as significantly increased AUC values compared to wildtype consistent with the presence of mechanical myotonia (Fig. 3f inset, p < 0.0001). Mbnl1^−/−^ ; ClC-1^ΔE7a^ force traces swiftly returned to baseline following tetanic stimulation with AUC values equivalent to wildtype, indicating the complete absence of mechanical myotonia. Continued firing of action potentials after tetani were observed in the Mbnl1^−/−^ muscles, but no activity following the stimulation artifact were seen in Mbnl1^−/−^ ; ClC-1^ΔE7a^ muscles (Fig. 3f lower panel). To better appreciate the differences in the EMG signal amplitudes and patterns, demodulation with a linear envelope approach was completed. There was a clear delay in return for to baseline for the Mbnl1^−/−^ recordings and swift return to baseline for Mbnl1^−/−^; ClC-1^ΔE7a^ recordings, indicating the presence and absence of electrical myotonia, respectively (Fig. 3g). Overall, the recordings of ClC-1 currents and the paired *in vivo* force-contraction/EMG experiments indicate that the gene-based exclusion of E7a was successful in both recovering ClC-1 channel function to wildtype-levels in Mbnl1^−/−^ mice, and that this was successful in completely eliminating myotonia from these mice, as previously accomplished by splice-switching ASOs^8,12^.

### Absence of myotonia improves specific-force production in *soleus* muscle

The impact of myotonia on muscle performance in Mbnl1^−/−^ mice was assessed by *ex vivo* force-contraction recordings in *soleus* muscles isolated from adult (4-6-month-old) WT, ClC-1^ΔE7a,^ Mbnl1^−/−^, and Mbnl1^−/−^ ; ClC-1^ΔE7a^ mice. Muscles were subjected to various stimulation protocols, including isolated 150Hz tetanic stimuli (500ms) designed for maximal activation, as shown by representative traces in Figure 4a. Peak specific force generated by Mbnl1^−/−^ muscles (144.9 ± 6.7 mN/mm^2^) was approximately 28% less than WT (201.3 ± 7.0 mN/mm^2^; p < 0.0004) with a partial recovery in force seen in the Mbnl1^−/−^ ; ClC-1^ΔE7a^ samples (178.0 ± 9.3 mN/mm^2^); Mbnl1^−/−^ ; ClC-1^ΔE7a^ recordings were not significantly different than WT (p = 0.2871). We observed no weakness in the ClC-1^ΔE7a^ mice alone (p = 0.8087; Fig. 4b). Similar trends in specific force output were seen across stimulation frequencies, with a generalized reduction in Mbnl1^−/−^ muscle compared to WT that was partially recovered in Mbnl1^−/−^ ; ClC-1^ΔE7a^ muscle (Fig. 4c). However, when force output for each sample was normalized, their peak tetanic force produced across stimulation frequencies there is a rightward shift in force-frequency for Mbnl1^−/−^ recordings, indicating a decreased sensitivity to stimulation frequency (i.e., an increased simulation frequency is required to generate the same fraction of specific force for a given muscle; Fig. 4d). This was quantified by the frequency required for half maximal stimulation, which was significantly larger in Mbnl1^−/−^ muscle compared to all other groups tested (Fig. 4e). Interestingly, when the values for cross-sectional area (CSA) of each of the soleus muscles used to collect force-contraction recordings from was inspected, there was an approximately 26% increase in CSA of Mbnl1^−/−^ solei compared to WT (p-adj. = 0.0288). This change in CSA was not seen in Mbnl1^−/−^ ; ClC-1^ΔE7a^ muscles, which were not significantly different from WT (p-adj. = 0.9466; Fig. 4f).

**Figure 4.**
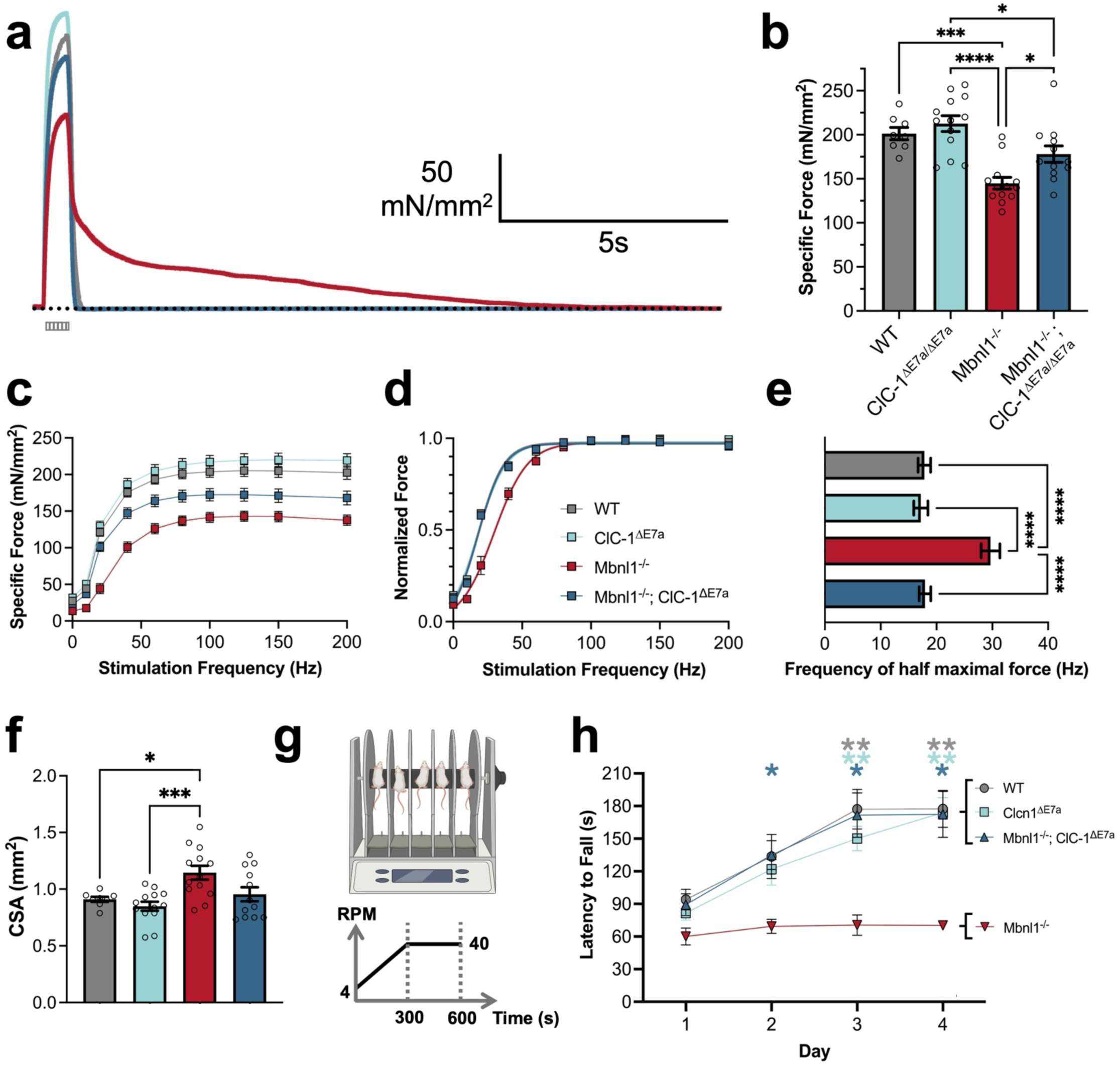
Non-myotonic Mbnl1 knockout mice produce increased contractile force compared to myotonic mice. *In vitro* force-contraction of soleus muscles isolated from 4-6-month-old WT, ClC-1^ΔE7a/ΔE7a,^ Mbnl1^−/−^, and Mbnl1^−/−^ ; ClC-1^ΔE7a/ΔE7a^ mice. (a) Representative specific force traces in response to a tetanic stimulus (150 Hz; 500 ms), scale as shown. Genotypes represented by colors consistent with the shading of bars in panel b. (b) Peak specific force generated during tetanic stimulation (150 Hz, 500 ms). Data plotted mean ± SE with individual replicates overlayed as dots. (c) Specific force versus frequency graph based on peak force generated for tetanic stimuli (500 ms) at stimulation frequencies ranging from 1Hz to 200Hz. Data presented mean ± SE with connecting lines. (d) Force normalized to peak tetanic force generated across all stimulation frequencies plotted versus stimulation frequencies. Data presented mean ± SE with line representing average fit of the data to a Boltzmann sigmoidal curve. (e) Frequency of half maximal force generation as determined by the Boltzmann sigmoidal fits in panel d. Data presented mean ± SE. (f) Cross-sectional areas (CSA) of soleus muscles isolated for force contraction. Data plotted mean ± SE with individual replicates overlayed as dots. (g) To assess in overall muscle function, rotarod experiments with a linear acceleration protocol were completed, as previously utilized for the assessment of DM1 mouse models. (h) Mice were run on the rotarod for 4 consecutive days (4 trials/mouse/day; 5-6 mice/genotype) with the latency to fall during the linear acceleration protocol assessed. Mean ± SE for WT, ClC-1^ΔE7a/ΔE7a,^ Mbnl1^−/−^, and Mbnl1^−/−^ ; ClC-1^ΔE7a/ΔE7a^ mice plotted across all the experimental days and analyzed with a repeated measures two-way ANOVA with Geisser-Greenhouse correction with Tukey’s multiple comparisons to assess the effects of experimental day and genotype on latency to fall. The interaction between genotype and experimental day was determined to be significant (F (9, 57) = 4.020, p = 0.005). Simple main effects of genotype were significant (p=0.0002) and main effects of experimental day were also found to be significant (p<0.0001). WT, ClC-1^ΔE7a,^ and Mbnl1^−/−^ ; ClC-1^ΔE7a^ mice all featured significant (p<0.05) improvements in performance on the final experimental day (day 4) as compared to the first day (day 1), whereas Mbnl1^−/−^ did not display any significant change in their performance (not shown). The colored stars above each trial day indicate the multiplicity adjusted p-value comparing for comparisons of Mbnl1^−/−^ versus each of the other respective genotypes (as per color). Other statistical comparisons in panels b, e, and f represent one-way ANOVA tests with Tukey’s post-hoc test. Multiplicity adjusted p-values shown for significant comparisons, as p-value < 0.05 *, 0.01 **, 0.001 ***, and 0.0001 ****.

### Absence of myotonia improved rotarod performance in Mbnl1 KO mice

Overall *in vivo* muscle function was assessed by rotarod with a linear acceleration protocol, as previously used to study mouse models of myotonic dystrophy (Fig. 4g)^19^. Across the four days of trials, WT, ClC-1^ΔE7a,^ and Mbnl1^−/−^ ; ClC-1^ΔE7a^ improved in the latency to fall. However, the Mbnl1^−/−^ mice failed to improve featuring a latency to fall significantly shorter than WT, ClC-1^ΔE7a,^ and Mbnl1^−/−^ ; ClC-1^ΔE7a^ mice (Fig. 4h).

### Absence of myotonia reduced histopathological changes seen in muscle of DM1 mice

Skeletal muscle in DM1 mouse models exhibit numerous histopathologic changes similar to those seen in DM1 patients, including increased central nucleation and alterations in fiber type compositon^8,13,20^. Therefore, to assess the impact of myotonia on the histopathologic hallmarks of DM1, we completed hematoxylin and eosin (H&E) staining to assess central nucleation and immunohistochemistry to assess fiber type distributions as well as myofiber morphometry across various hindlimb muscles and diaphragm strips from 4-6-month-old mice. Here, the *tibialis anterior* (TA) muscle is shown with representative H&E-stained transverse sections, Figure 5a. We observed a significant increase in the percent of fibers with central nuclei present in Mbnl1^−/−^ muscles, characteristic of muscle samples collected from both DM1 mouse models and patients^21,22^. Notably, Mbnl1^−/−^ ; ClC-1^ΔE7a^ muscle displayed a significantly reduced percent of fibers with central nuclei (∼56% less) as compared to Mbnl1^−/−^ samples (p = 0.0004, Fig. 5b), which indicates that myotonia may play a role in driving central nucleation in DM1 mice. Similar trends were observed in the *quadriceps* muscle (Fig. S6a,b). Immunohistochemistry with antibodies targeting MyHC isoforms for fiber-typing and laminin for fiber morphometry was completed with representative TA transverse sections shown in Figure 5c. In Mbnl1^−/−^ samples, we observed a significant increase in fibers staining positive for the type 2a MyHC isoform, with a concomitant decrease fiber positive for the type 2b MyHC isoform compared to WT. This shift in fiber type distribution was not observed in the Mbnl1^−/−^ ; ClC-1^ΔE7a^ samples. Furthermore, no shifts in the percentage of type 1 or type 2x fibers were observed across any of the groups (Fig. 5d). Overall, these changes remain consistent across the other hindlimb muscles tested, all of which have been shown to exhibit myotonia in the Mbnl1 knockout model of DM1^21^. Interestingly, there were not shifts in fiber type distribution seen in the diaphragm, which was subsequently confirmed to lack myotonia in Mbnl1^−/−^ mice and aligns with recent studies in myotonia congenita (Fig. S6c,d,e, S7a)^23^. We also observed a significant increase in the Feret’s minimum diameter in the Mbnl1^−/−^ TA, whereas Mbnl1^−/−^ ; ClC-1^ΔE7a^ samples were similar to WT (Fig. 5e). When muscle fiber morphometry was analyzed in specific fiber-types, it was shown that the increase in the Feret’s minimum diameter was predominantly seen in the type 2 subtypes (Fig. S7b). Additionally, ClC-1^ΔE7a^ control samples were near WT all parameters assessed, including percent of fibers with central nuclei, fiber-type distributions, and fiber-morphometry, across all muscle groups tested (Fig. 5, S6, S7).

**Figure 5.**
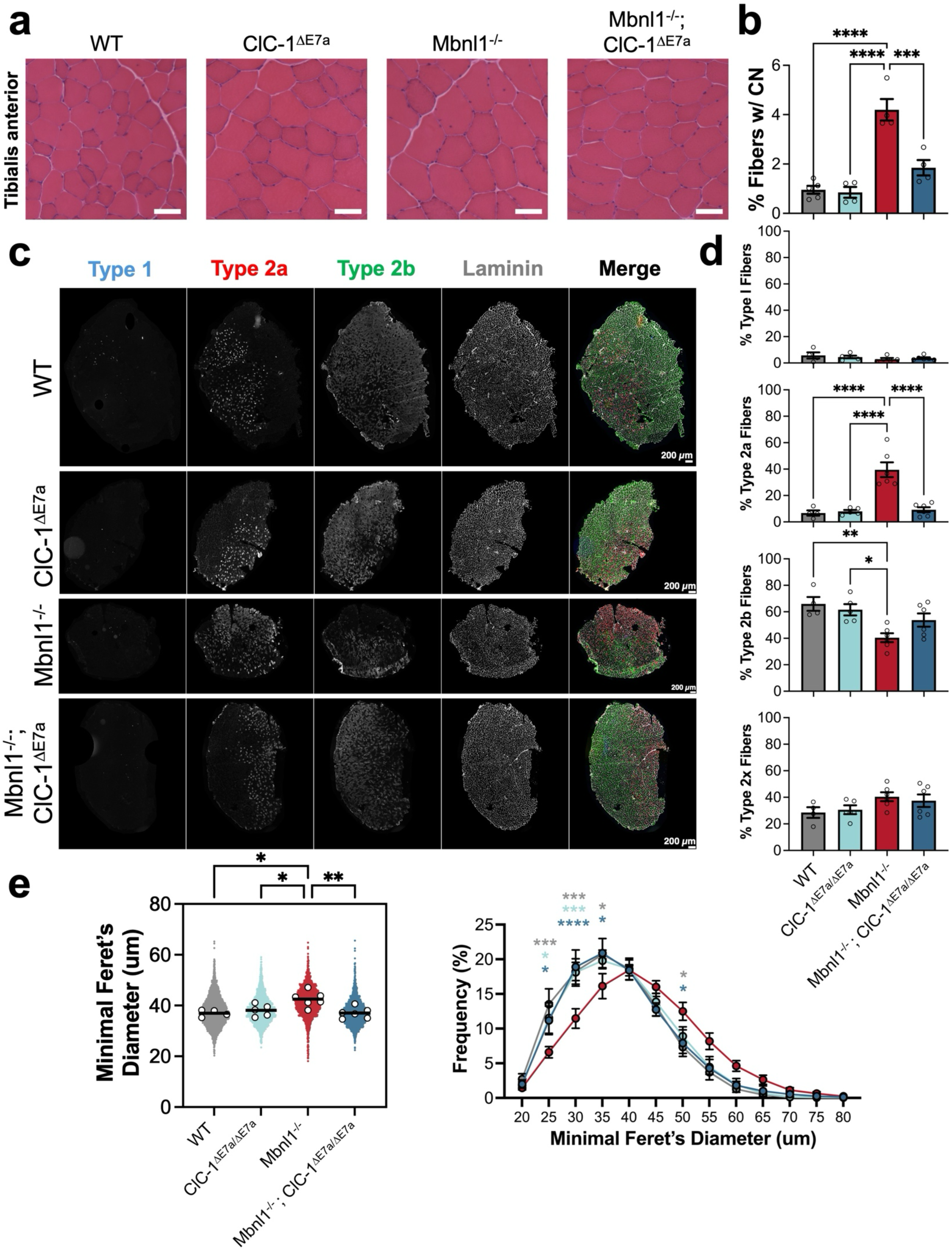
Myotonia is a driver of histopathologic changes in Mbnl1 knockout skeletal muscle. (a) Representative images of tibialis anterior transverse sections stained with hematoxylin and eosin (H&E) from WT, WT, ClC-1^ΔE7a/ΔE7a,^ Mbnl1^−/−^, and Mbnl1^−/−^ ; ClC-1^ΔE7a/ΔE7a^ mice (4-6-month-old). Scale bars shown represent 50 micrometers. (b) Percent of fibers with central nuclei present on H&E strained tibialis anterior samples (n = 4-5 mice/genotype). (c) Fiber-typing by myosin heavy chain (MyHC) isoform was completed by immunohistochemistry, representative images shown, with each MyHC isoform channel and laminin counterstain shown in grayscale and with the merged image pseudo-colored (based on color of channel labels). (d) Percent of fibers staining for each of the respective MyHC isoforms (n = 4-6 mice/genotype). (e) Fiber morphometry was completed based on laminin staining with minimum Feret’s diameter determined for each fiber plotted as a scatter plot (left) and histogram (right). For the scatter plot, small dots represent aggregation of all fibers analyzed across biological replicates, whereas larger dots filled in white represent mean of each biological replicate. Statistical comparisons in (b), (d), and (e, left) with one-way ANOVAs with Tukey’s multiple comparisons. Multiplicity adjusted p-values of significant comparisons presented as * p<0.05, ** p<0.01, *** p<0.001, and **** p<0.0001. Comparisons in (e, right) represent a RM two-way ANOVA assessing the effects of genotype and minimal Feret’s diameter on the frequency distribution with Tukey’s multiple comparisons. There was a significant interaction between genotype and minimal Feret’s diameter (F (36, 204) = 2.442, p <0.0001). Additionally, the main effects of Feret’s minimum bin (p<0.0001) and genotype (p<0.0001) were found to be significant. Stars above both histograms represent multiplicity adjusted p-values of significant comparisons between the genotype associated with the star color and Mbnl1^−/−^ mice for that bin, as no comparisons between the other groups were found to be significant: * p<0.05, *** p<0.001, ****, p<0.0001. Data presented as mean ± SEM with individual values.

**Figure 6.**
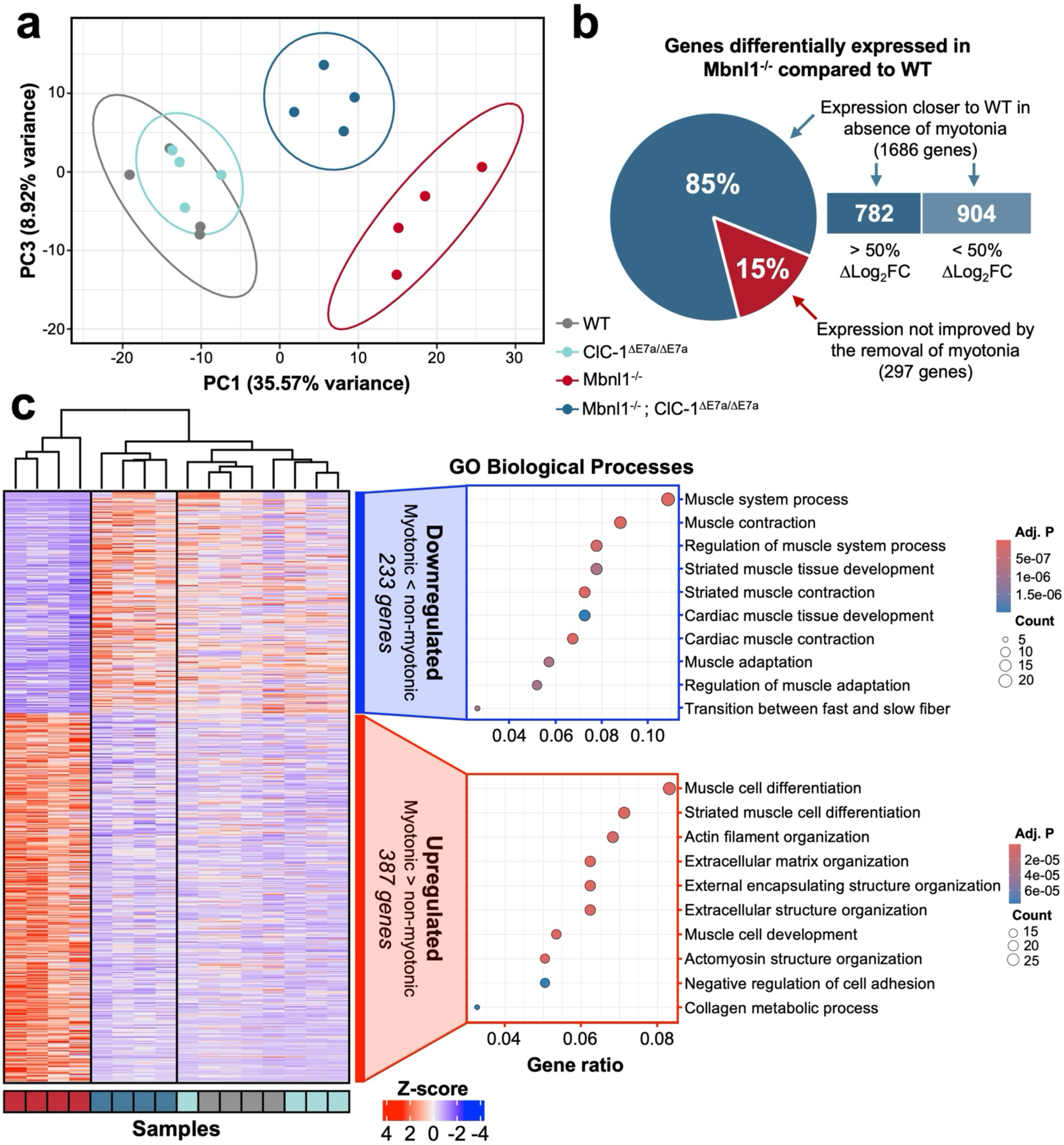
Myotonia alters transcript expression in Mbnl1−/− mice. RNA sequencing (conventional deep, short-read with paired-ends) was completed on quadriceps muscle isolated from 4-6-month-old WT, ClC-1^ΔE7a/ΔE7a,^ Mbnl1^−/−^, and Mbnl1^−/−^ ; ClC-1^ΔE7a/ΔE7a^ mice (n = 4 mice/genotype) to assess the impact of myotonia on the transcriptomic changes seen in Mbnl1^−/−^ mice. (a) Principal component analysis was completed for all samples based on the top 500 genes with the greatest variance with samples plotted against PC1 and PC3 (PC2 excluded as top loading genes represented predominantly those localized to sex chromosomes). Ellipses represent 95% confidence boundary of sample clusters. (b) Differential expression analysis was completed using DESeq2 with pairwise comparisons between Mbnl1^−/−^ and WT as well as Mbnl1^−/−^ ; ClC-1^ΔE7a/ΔE7a^ and WT to allow for the determination of the genes with expression changes in Mbnl1^−/−^ mice as well as the impact of the absence of myotonia in Mbnl1^−/−^ ; ClC-1^ΔE7a/ΔE7a^ on the expression of those genes, respectively. There were 1,983 genes found to be dysregulated in Mbnl1^−/−^ mice (⎸log_2_FC ⎸> 0.5 and adj. p-value < 0.05); in Mbnl1^−/−^ ; ClC-1^ΔE7a/ΔE7a^ the expression of 1,686 of these genes was closer to wildtype indicated by a log_2_FC versus WT closer to 0. For 782 of the 1,686 genes, the log_2_FC of Mbnl1^−/−^ ; ClC-1^ΔE7a/ΔE7a^ versus WT was half that or less compared to the log_2_FC of Mbnl1^−/−^ versus WT. (c) Direct comparison of the differential expression between Mbnl1^−/−^, and Mbnl1^−/−^ ; ClC-1^ΔE7a/ΔE7a^ mice yielded 620 differentially expressed genes with a heatmap of the normalized expression of these genes shown for all samples of the experimental and control groups. Based on the expression of these genes, unsupervised sorting of the samples was completed with the Mbnl1^−/−^ and Mbnl1^−/−^ ; ClC-1^ΔE7a/ΔE7a^ clustering in isolated groups and control samples (WT and ClC-1^ΔE7a/ΔE7a^) co-clustering. Gene ontology analysis of biological processes represented in the downregulated and upregulated genes in Mbnl1^−/−^ versus Mbnl1^−/−^ ; ClC-1^ΔE7a/ΔE7a^ samples was completed. Scales as shown.

**Figure 7.**
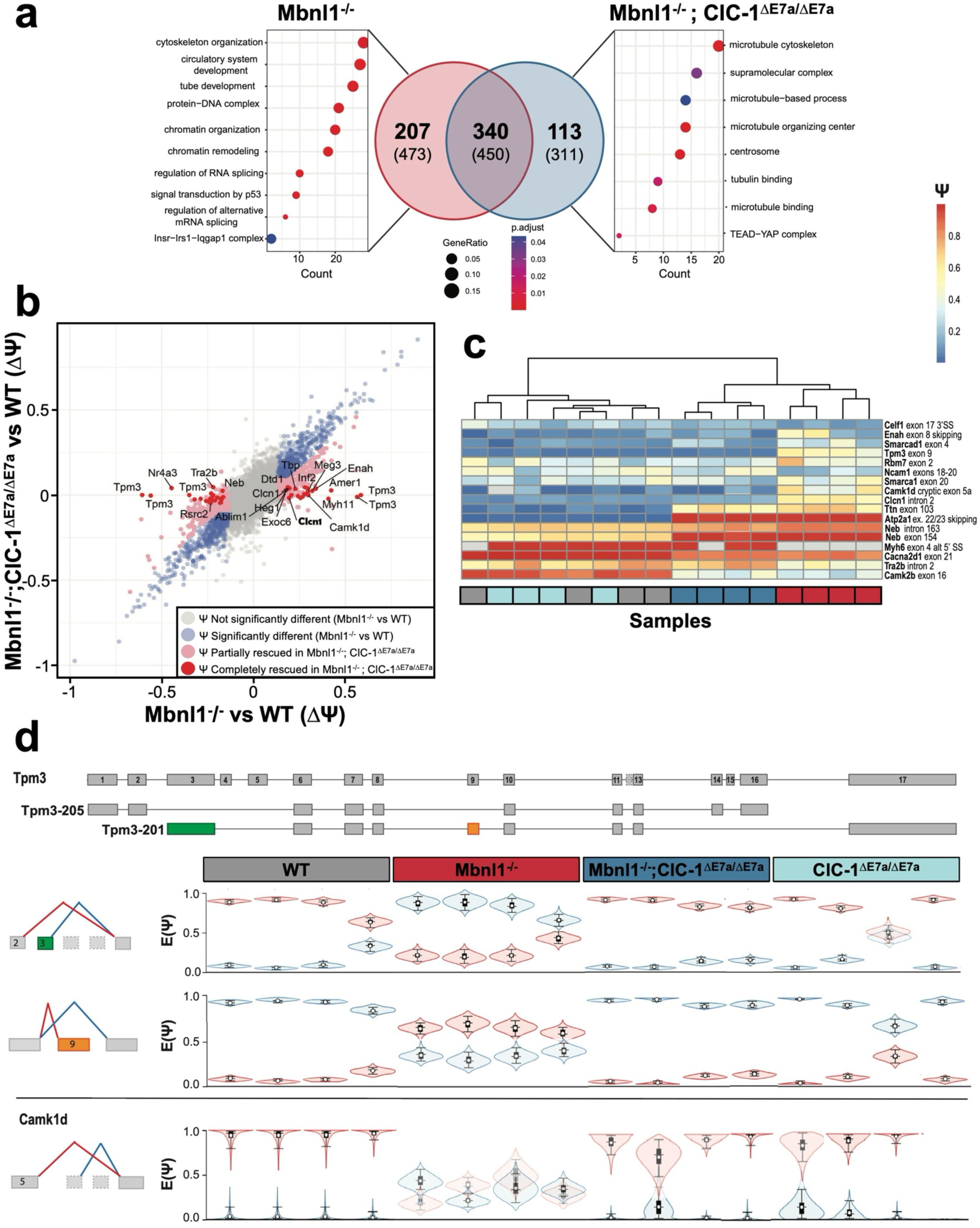
Myotonia regulates specific splicing events. (a) Overlap of genes mis-spliced in muscle of myotonic Mbnl1 KO mice (Mbnl1^−/−^) and Mbnl1 KO mice without myotonia (Mbnl1^−/−^ ; ClC-1^ΔE7a/ΔE7a^), both compared to WT mice. Most genes (52%, 340 genes) are commonly mis-spliced (FDR > 90%, delta-PSI > 0.1 & < −0.1), while there are 207 and 113 genes uniquely mis-spliced in myotonic and non-myotonic Mbnl1 KO muscle, respectively (note that unique genes might have the same event in both conditions, but miss the threshold set for delta-PSI to be significantly different). Unique genes were analyzed regarding biological pathway (dot plot, top 10 GO terms), which revealed that genes that are only mis-spliced in myotonic Mbnl1KO mice are involved in cytoskeletal and chromatin organization as well as RNA splicing, whereas in non-myotonic Mbnl1 KO mice, unique genes fall mainly in cytoskeletal organization. (b) Splicing events in Mbnl1^−/−^ and Mbnl1^−/−^;ClC-1^ΔE7a/ΔE7a^ were each compared to WT mice and the PSI-values were plotted as a scatter plot (each dot represents one ASE with some transcripts having multiple ASE). Events that are not changing between Mbnl1^−/−^ and WT are displayed in grey, while events with PSI-value above 0.1 and below −0.1 and an FDR above 80% are shown in blue. Pink events are partial rescues defined as having more than a 0.15 PSI shift in Mbnl1^−/−^ ; ClC-1^ΔE7a/ΔE7a^ towards control. Events with a complete rescue (changing in Mbnl1^−/−^ vs WT and non-changing in Mbnl1^−/−^;ClC-1^ΔE7a/ΔE7a^ vs WT at an FDR of > 95%) in mice without myotonia are shown in red and are labeled. The positive control Clcn1 exon7 skipping is marked in bold. (c) Heatmap of selected genes ranging from complete rescue to unaffected. Rows represent individual splice events (labels right) and columns represent biological replicates, arranged by unsupervised sorting (genotype denoted by colored box below). (d) Tpm3 shows the strongest rescue in mice without myotonia. In Mbnl1^−/−^ mice, exon 3 and exon 9 are included with a PSI-value of +0.65 and 0.7, respectively. The isoform including exons 3 and 9 is Tpm3-201, in which exon 3 becomes the first exon, while the exon 5 codes for exon 5. In wildtype mice however, exons 1,2, 10, 14 and 15 which belong to isoform Tpm3-205 (Fig. S8).

### Myotonia drives changes in transcript expression in skeletal muscle of Mbnl1 knockout mice

DM1 is predominantly characterized as disease of altered transcript splicing; however, derangements in the expression of numerous genes have been observed both in DM1 patients and mouse models of DM1^9,24–26^. Efforts to understand the cause of changes in transcript expression in DM1 led to numerous hypotheses ranging from direct interaction of transcripts with toxic RNA repeats to RNA interference^9,27^. However, comparisons between mouse models of myotonia congenita and DM1 have shown similar trends in transcript expression changes for almost one-fourth the genes differentially expressed in DM1 mouse models^9^. Here, we were able to directly compare changes in transcript expression between myotonic (Mbnl1^−/−^) and non-myotonic (Mbnl1^−/−^ ; ClC-1^ΔE7a/ΔE7a^) DM1 model mice with deep, short-read RNA-seq (150bp paired-end reads; 66.25 ± 8.26 million unique reads (mean ± IQR) mapped to genes with at least 50 supporting reads across all samples) on RNA isolated from quadriceps muscle from 4-6-month-old WT, ClC-1^ΔE7a/ΔE7a,^ Mbnl1^−/−^, and Mbnl1^−/−^ ; ClC-1^ΔE7a/ΔE7a^ mice (n=4 mice/genotype). Differential expression analysis was completed with DESeq2, and pair-wise contrasts were completed for all permutations of the four genotypes ^28^. Moreover, normalized counts of manipulated genes (*Mbnl1* and *Clcn1*) were inspected to provide internal validity of sequencing results (Fig. S8a,b). Principal component analysis (PCA) using the top 500 genes with the most variance across all samples revealed co-clustering of WT and ClC-1^ΔE7a/ΔE7a^ (circle represent 95% CI) control samples as well as tight clustering of the Mbnl1^−/−^ and Mbnl1^−/−^ ; ClC-1^ΔE7a/ΔE7a^ with dispersion of the groups particularly across PC1; Mbnl1^−/−^ samples were localized the furthest from the control samples with Mbnl1^−/−^ ; ClC-1^ΔE7a/ΔE7a^ intermediate, or more proximal to the control samples, along PC1 (Fig. 6a, Fig. S8c,d,e).This supports the hypothesis that myotonia may account for some of portion of the altered transcript expression seen in Mbnl1 knockout mice. Furthermore, when the genes with the highest loadings in PC1 were further interrogated by gene ontology analysis the biological pathway most represented was skeletal muscle contraction (Fig. S8f). Specifically, several genes involved in the contractile apparatus were mis-regulated, including several myosin light and heavy chain isoforms, Tnnt1, that encodes the slow skeletal muscle isoform of troponin T, and Tnni1 that slow-twitch skeletal muscle isoform of troponin I. With several other regulators of skeletal muscle contraction (e.g., Csrp3) that have been associated with skeletal muscle contractile function and myopathic diseases ^29–31^. Next, to directly assess the proportion of genes dysregulated in Mbnl1^−/−^ muscle that have expression levels closer to WT in the absence of myotonia (i.e., in Mbnl1^−/−^ ; ClC-1^ΔE7a/ΔE7a^ samples), we completed pair-wise differential expression analysis between Mbnl1^−/−^ and WT samples as well as Mbnl1^−/−^ ; ClC-1^ΔE7a/ΔE7a^ and WT samples, then compared the resulting gene sets. Overall, we found 1,983 genes mis-regulated in Mbnl1^−/−^ samples compared to WT (∣log_2_FC∣ > 0.5, p-adj. < 0.05). For 1,686 of these genes (85%), the log_2_FC for the Mbnl1^−/−^ ; ClC-1^ΔE7a/ΔE7a^ samples compared to WT was closer to zero, with 782 of these genes with a greater than 50% change in log_2_FC (Fig. 6b). Together, this indicates that myotonia plays the predominant role in driving expression level changes for more than 39% of transcripts in Mbnl1^−/−^ muscle. To provide a better understanding of the specific transcripts with differential expression in myotonic and non-myotonic Mbnl1^−/−^ mice, we then directly compared Mbnl1^−/−^ to Mbnl1^−/−^ ; ClC-1^ΔE7a/ΔE7a^ samples and found 620 differentially expressed genes (∣log_2_FC∣ > 0.5, p-adj. < 0.05) between Mbnl1^−/−^ and Mbnl1^−/−^ ; ClC-1^ΔE7a/ΔE7a^. Gene ontology (GO) analysis was completed on the up- and down-regulated genes, with biological processes shown as dot plots in Figure 6c. The top biological process represented in the subset of genes downregulated in Mbnl1^−/−^ muscle compared to Mbnl1^−/−^ ; ClC-1^ΔE7a/ΔE7a^ were related to skeletal muscle contraction and muscle system processes, including several myosin light heavy chain isoforms (*Myh6*, *Myh7*, *Myh7b*, *Myl2*, *Myl6b*), genes encoding proteins in the contractile apparatus (*Tnnc1*, *Tnnt1*, *Tnni1*, *Tpm3*), as well as several genes involved in calcium signaling (*Atp2a2*, *Strit1*, *Ryr2*, *Calm1*, *Calm3*). Whereas the biological process most upregulated in Mbnl1^−/−^ compared to Mbnl1^−/−^ ; ClC-1^ΔE7a/ΔE7a^ samples was for skeletal muscle differentiation, including *Cacnb4*, genes involved in cellular signaling such as *Camk2d*, *Myf6*, and *Popdc3*, and several genes encoding proteins important for skeletal muscle cytoskeletal and sarcomere structure such as *Flnc*, *Myom2*, *Col14a1*, and *Tmod2*. Similarly, gene ontology analysis was also completed for molecular functions with dysregulation of genes encoding many ion channels observed, indicating possible compensatory regulation of skeletal muscle excitability in the setting of the loss of the main resting inhibitory conductance on the sarcolemma. Overall, this exploratory RNA-seq analysis suggests that myotonia plays an important role in the mis-regulation of transcript expression seen in Mbnl1^−/−^ mice with possible implications in progresses related to skeletal muscle function and differentiation.

### Removal of myotonia reverses mis-splicing of a specific set of genes

To investigate whether myotonia influences alternative splicing, we analyzed all samples using the splicing analysis tool MAJIQ, which detects both simple (e.g., exon skipping, intron retention) and complex splicing events (e.g., triple exon skipping versus intron retention). For a comprehensive overview, we compared Mbnl1^−/−^ and Mbnl1^−/−^ ; ClC-1^ΔE7a^ mice to wild-type mice separately.

MAJIQ identified a higher number of splicing events in Mbnl1^−/−^ mice (923 events across 547 genes) compared to Mbnl1^−/−^ ; ClC-1^ΔE7a^ mice (761 events across 453 genes), applying thresholds of ΔPSI > 0.1 or < −0.1 with a false discovery rate (FDR) < 0.05. This observation suggests that myotonia may impact splicing regulation. Comparative analysis of the affected genes and splicing events (Fig. 7a) revealed that 52% of the genes were shared between both conditions. Notably, Mbnl1^−/−^ mice exhibited a greater number of unique splicing events (473) compared to Mbnl1^−/−^ ; ClC-1^ΔE7a^ mice, which displayed 311 unique events across 113 genes. This finding indicates that myotonia might not only contribute to the occurrence of specific mis-splicing events, but could also mitigate certain splicing anomalies induced by Mbnl1 depletion.

Interestingly, several splicing factors including *Srsf6*, *Prpf39*, *Rbm7*, *Tra2b*, *Rbfox2*, and *Hnrnpu*, exhibited alternative splicing in Mbnl1^−/−^ mice with myotonia, but not in those without. Consistently, pathway analysis of uniquely mis-spliced genes in both conditions highlighted the regulation of RNA splicing as one of the ten most significantly mis-regulated pathways (Fig. 7a, left dot plot), alongside pathways involved in cytoskeleton and chromatin organization in Mbnl1^−/−^ mice (Fig. S9).

Chromatin remodeling is particularly noteworthy, as mechanical stress from muscle contraction has been shown to profoundly influence chromatin architecture^32–34^. Given that myotonia leads to prolonged aftercontractions, this may have implications for chromatin organization^35^. In Mbnl1^−/−^ mice without myotonia, uniquely mis-spliced genes are predominantly associated with cytoskeleton organization and the Yap-Tead complex, comprising the transcription factor YAP1 and its coactivator TEAD1, both of which respond to mechanical stimuli (Fig. 7a, right dot plot). Notably, exon skipping events in Yap1 and Tead1 were detected exclusively in mice without myotonia, absent in those with myotonia.

### Tropomyosin 3 mis-splicing is fully rescued in Mbnl1^−/−^ ; ClC-1^ΔE7a/ΔE7a^ mice

We conducted a detailed comparison of alternative splicing events between Mbnl1^−/−^ and Mbnl1^−/−^ ; ClC-1^ΔE7a^ mice, confirming a strong correlation for the majority of shared events (Fig. 7b, blue). Additionally, we identified numerous partial rescue events in non-myotonic mice, characterized by a shift toward control levels of at least 0.15 dPSI (shown in pink, FDR < 0.2). Complete rescue events were defined as cases where splicing in non-myotonic mice remained unchanged compared to wild type, while Mbnl1^−/−^ mice exhibited a shift of at least ± 0.1 dPSI (shown in red, FDR < 0.05). Notably, *Clcn1* exon 7 was correctly classified as ‘rescued.’ Strikingly, this analysis revealed that all alternative splicing events detected in tropomyosin 3 (Tpm3) in Mbnl1^−/−^ mice were fully rescued in Mbnl1^−/−^ ; ClC-1^ΔE7a^ mice. Similarly, the genes *Enah* and *Ablim1*, which are frequently mis-spliced in DM1 patients^36,37^, displayed complete rescues of at least one detected splicing event. Correlating myotonia severity with the mis-splicing of these genes in patients would be particularly insightful. The heatmap in Figure 7c illustrates PSI values for selected splicing events that remain unchanged between myotonic and non-myotonic mice (*Atp2a1, Neb, Ttn, Cacna2d1, Camk2b*), as well as those that exhibit partial or complete rescue in at least two animals. Unsupervised clustering further supports the idea that non-myotonic mice represent an intermediate state between Mbnl1^−/−^ and WT mice.

Among the rescued events, *Tpm3* emerged as the most prominent transcript, warranting a closer examination. MAJIQ analysis identified five splicing events that differed between Mbnl1^−/−^ and WT mice, with high dPSI values (up to 0.7) and strong confidence (FDR < 0.05), two of which are shown in Figure 7d. In myotonic Mbnl1^−/−^ mice, *Tpm3* exons 3 and 9 were preferentially included, whereas exons 1, 2, and 14 were predominantly used in WT mice (Fig. 7d). These exon distributions correspond to two distinct *Tpm3* isoforms, Tpm-201 and Tpm-205.

## Discussion

We demonstrated that CRISPR-mediated excision of the E7a sequence from the *Clcn1* gene was sufficient to restore ClC-1 channel function and eliminate myotonia from the Mbnl1^−/−^ model of DM1 with a resultant decrease in the overt manifestations of myopathy, including and maybe most importantly, significantly improved force production. Accordingly, this novel model shows the benefits of ideal, lifelong anti-myotonic therapy. Moreover, we also presented (i) the first use of targeted long-read sequencing in the field of myotonic dystrophy, to our knowledge, to assess Mbnl1-dependent transcript isoform changes for a gene with complex splice patterning, (ii) a novel method for the assessment of mechanical and electrical myotonia in DM1 mouse models, and (iii) a comprehensive analysis of the histopathological and transcriptomic impact of myotonia in skeletal muscle of DM1 mouse models.

Targeted long-read (third-generation) sequencing is emerging as useful technique for probing RNA processing and unique transcript isoforms for genes with complex splice patterning^38^. In the field of neuromuscular diseases, specifically in DM1, third-generation sequencing has primarily been used to provide improved diagnostic capabilities and the characterization of the toxic-repeat sequence; for example, efforts to better understand interruptions in the repetitive sequence^39,40^. Here, we leveraged long-read sequencing to better characterize changes in *Clcn1* transcript isoform utilization for in the absence of functional Mbnl1. Over the past decades since the implication of ClC-1 deficiency as the cause of myotonia in DM1, it has been shown that numerous splice events in *Clcn1* transcript, in addition to E7a, were upregulated in DM1 patients and disease models, including intron 2 retention^3^. However, these findings conflict with functional data, shown here with CRISPR-based gene editing (Fig. 3) or previously with splice-shifting E7a splicing specific ASOs^12^, that were capable of restoring WT or even supr-maximal levels of ClC-1 function in DM1 mouse models. This prompted the question, are these other splicing events regulated by Mbnl1 or is their upregulation the byproduct of changes in the overall stability of *Clcn1* transcripts?

In this study, and in agreement with previous studies, we demonstrate that loss of Mbnl1 activity leads to decreased steady-state *Clcn1* mRNA. This is thought to be driven largely by decreased mRNA stability stemming from the aberrant inclusion of E7a, causing a frameshift, and ultimately NMD of transcripts^4^. Accordingly, this leads to an overall reduction in the number of *Clcn1* transcripts that are not targeted by NMD processes, and thus other rarer aberrant splicing events that emanate from the spliceosome become relatively more abundant. Moreover, the presence of a second NMD causing splicing change (e.g., the simultaneous intron 2 retention and E7a inclusion in a single transcript), likely does not significantly alter the stability of the transcript more than a single NMD causing splice-event. This was supported by both the large increase in intron 2 retention seen in ADR samples, in which all transcripts possess at least one NMD-yielding event, as well as the relatively more abundant intron 2 retention in E7a containing transcripts in Mbnl1^−/−^ samples (Fig. S3d, S3e). Overall, these results support the conclusion that E7a inclusion is the primary event regulated by Mbnl1 in *Clcn1* transcripts, with the other events being the byproduct of changes in overall transcript stability. Furthermore, this raises the importance of sequencing transcripts with complex splicing patterning end-to-end, rather than individual splice events in isolation, when interrogating the effects of splice-shifting therapeutics.

Myotonia remains a key symptom and phenotype used to assess the severity of the DM1 disease state as well as evaluate the success of novel therapeutics in patients and disease models^41–44^. However, many commonly used assays lack methods of objective quantification, longitudinal monitoring, and do not fully assess both mechanical and electrical myotonia despite many efforts to develop assays that reliably quantify the severity of myotonia^45^. Here, we utilized both *in vivo* plantarflexion and paired posterior compartment subcutaneous EMG to provide methods for quantifying both mechanical and electrical force production. Despite its relatively undemanding utilization in this study (i.e., the detection of the presence or absence of myotonia), this technique will provide a platform for further studies tracking responses to therapeutic interventions as well as potentially provide further insight into emerging pathophysiological phenomenon, such as silent myotonia and sustained after-depolarizations, in myotonic disorders^46^.

Furthermore, through deep RNA-seq we were able to directly assess the impact of myotonia on transcript expression and splicing in mice lacking functional Mbnl1. We observed significant changes in transcript abundance between myotonic and non-myotonic Mbnl1^−/−^ mice with as high as 85% of transcripts in non-myotonic Mbnl1^−/−^ mice having expression levels closer to wildtype than myotonic samples. This is far greater than previously estimated (approximately 25%) through comparisons with mouse models of non-dystrophic myotonia^9^. Moreover, of the genes that featured significantly different expression levels between myotonic and non-myotonic Mbnl1^−/−^ samples, the biological pathways most significantly enriched were muscle contraction and differentiation, further substantiating the role of myotonia in driving muscle weakness. Interestingly, another molecular function that featured substantial differences was in other ion channels on the sarcolemma and transverse tubule membrane, including several potassium channels and their subunits (Kcnab1/Kcnh7/Kcnk3/Kcng4/Kcnb1/Kcnk2). This indicates the potential role of compensatory changes in the expression of other ion channels to modulate myotonia in DM1. These findings are important for consideration when studying alternative targets for the treatment of myotonic disorders, such as previously completed with the use of potassium channel agonists in myotonia congenita mouse models^47–49^. We also observed changes in transcript splicing for several in the setting of or absence of myotonia in Mbnl1^−/−^ mice, including tropomyosin isoforms, indicating a potential role of myotonia for regulating splicing changes in DM1. The mechanism for this remains to be elucidated however, as these experiments were completed in mice deficient in functional Mbnl1, absent of toxic-CUG repeats, these findings represent a process independent of functional Mbnl1. Notably, broader mis-regulation of splicing factors has been demonstrated in primary muscle cells isolated from people with DM1^50^ and in Mbnl1^−/−^ mice with, but not in those without, myotonia.

In this study, myotonia was found to play a previously undescribed role in orchestrating changes in splicing for several transcripts in DM1 mouse models. Tropomyosins are two-stranded, α-helical coiled-coil proteins that polymerize head-to-tail along actin filaments, regulating actin stability and function (reviewed in^51^). Each *Tpm* gene (*Tpm1-4*) encodes multiple isoforms with distinct expression profiles, localization, and functions^52^. Specifically, *Tpm* genes produce both high-molecular-weight (HMW) and low-molecular-weight (LMW) isoforms, where HMW isoforms serve as muscle-specific variants, while LMW isoforms function primarily in non-muscle cells. *Tpm3* is the predominant slow muscle fiber isoform among tropomyosins, with *Tpm-205* being the major HMW isoform critical for muscle contraction, whereas LMW *Tpm3* isoforms regulate T-tubule function and glucose uptake^53,54^. Interestingly, in Mbnl1^−/−^ mice, *Tpm3* is not only mis-spliced but also significantly downregulated (−2.7 log_2_FC). In Mbnl1^−/−^ ; ClC-1^ΔE7a^ mice, both splicing and transcriptional expression of *Tpm3* were largely restored (−0.2 log_2_FC). Despite clear ties between cellular excitability and transcript expression in cells such as neurons through the process known as, excitation-transcription coupling ^55^, the mechanism by which cellular excitability exerts changes in splice patterning remains unknown and thus warrants further studies into the process of excitation-splicing coupling.

Overall, the work presented here lays the foundation for establishing a pathologic role of myotonia in the development of myopathy in myotonic dystrophy (type 1 and 2) and provide initial indications that long-term removal of myotonia from myotonic dystrophy may prevent some of the progressive myopathy seen in the disease. Accordingly, further pre-clinical studies utilizing clinically available anti-myotonic agents or even clinical trials with longer-term follow-up are needed to fully ascertain whether these changes will hold true in settings more tangible to clinical practice^11^. Moreover, in the era of novel RNA targeted therapeutics designed to directly target the pathologic processes underlying DM1, there likely could still be an important role for these anti-myotonic treatments based on financial barriers as well as the potential for combinatorial therapy with additive or synergistic benefit.

## Methods

### Mouse lines

A transgenic mouse line that prevents the inclusion of the alternate exon 7a (E7a) in Clcn1 transcripts was generated with CRISPR/Cas9 designed for targeted excision of the E7a sequence from the *Clcn1* gene. Single guide RNAs (sgRNAs) targeted the intronic sequences flanking this alternative exon; this was to ensure complete removal of the splice donor/acceptor sites, as this has been previously shown to be successful to force the exclusion of a targeted exons while not perturbing other splice events in the transcripts^10,56^. Sequences of sgRNAs were 5’-GGAATCGACACAGTTCATGTGGG -3’ and 5’-GAACTTATGGGTGGGCAACCAGG-3’. The mouse line was generated on a C57BL/6J background and extensively backcrossed (>6 generations) with wild-type mice to eliminate possible off target mutations. Standard PCR with primers flanking the excised region (Supplemental Table 1a) were used for genotyping from genomic DNA and the forward primer was used for Sanger sequencing (Eurofins Scientific) to verify the correct excision from the *Clcn1* gene. Subsequently, this line was extensively backcrossed (>6 generations) with FVB/N mice, as the model Mbnl1^−/−^ used here are all maintained on a congenic FVB/N background. All other experiments included in this manuscript were completed on mice with a congenic FVB/N background. Homozygous mice lacking exon 7a from both *Clcn1* alleles are denoted as ClC-1^ΔE7a/ΔE7a^. For some behavioral and functional experiments both heterozygote (as ClC-1^ΔE7a/+^ and homozygote (as ClC-1^ΔE7a/ΔE7a^) mice were used, as the excision of only a single allele was shown to be successful in eliminating myotonia, this was denoted as ClC-1^ΔE7a^. The Mbnl1-null mouse that was used featured the targeted excision of exon 3 and was extensively characterized previously as a moderate DM1 model featuring skeletal muscle myotonia and increased central nuclei^13^. For simplicity, these mice are indicated as Mbnl1^−/−^. Additionally, for a Clcn1 non-sense mediated decay (NMD) control mouse model, *adr-mto2J* mice (ADR) were obtained from Jackson Laboratories (Bar Harbor, MA) and extensively backcrossed to the FVB/N background. These mice feature a frameshift mutation in exon 18 of the *Clcn1* gene that causes a premature termination codon in exon 19—previously shown to induce significant NMD^4^. Standard genotyping methods utilizing genomic DNA isolation and PCR with agarose gel electrophoresis (ClC-1^ΔE7a,^ Mbnl1^ΔE3^) or Sanger sequencing (ADR) were used to confirm all exon deletions and/or mutations (Supplemental Table 1a). Both male and female mice were used for all experiments, as no sex-based differences were shown with these strains in both the previous literature and preliminary experiments with these mice^13,57^. All experiments were completed as outlined by an animal protocol approved by the University of Rochester Committee on Animal Resources (UCAR; equivalent to IACUC). Animals were housed in the university’s vivarium in climate-controlled rooms with micro-isolation employed until the time of experiments.

### RNA isolation

RNA was isolated from quadriceps muscle excised from mice that was immediately frozen in liquid nitrogen and stored at −80°C. Approximately 50-100mg of tissue was homogenized using a bead mill (Bead Ruptor, Omni International) with ceramic beads (Fisherbrand) in TRI-reagent (Molecular Research Center) for initial RNA isolation. RNA was then purified (re-isolated) with RNeasy mini columns (Qiagen) with on-column DNase I-treatment (RNase-free DNase I, Qiagen), as previously described^58^. Yield and purity were assayed with a NanoDrop (Thermo Scientific), and only high-quality samples were used for RT-(q)PCR and sequencing.

### RT-(q)PCR

To assay transcript splicing as well as confirm E7a exclusion in the novel ClC-1^ΔE7a^ line, standard RT-PCR protocols with primers located in exons flanking alternatively spliced regions was utilized. Briefly, reverse transcription to create cDNA libraries from each muscle total RNA sample was completed using *Superscript II* (Invitrogen) primed with oligo(dT)_15_ (Promega) using the standard manufacture’s protocol with 1ug total RNA reverse transcribed per 20uL reaction. cDNA used for long-read sequencing (PacBio) were treated with RNase H following reverse transcription, discussed further in long-read sequencing methods. For targeted assessment of alternative splicing, PCR products were amplified with Phusion polymerase (NEB) with primers in exons flank alternative spliced regions (Supplemental Table 1a) and visualized using standard gel electrophoresis with 2% agarose gels stained with ethidium bromide (EtBr). Gels were imaged using either a GelDoc or ChemiDoc gel imaging system (BioRad) equipped with a UV tray. Images were processed with Image Lab (BioRad) and band densitometry was determined with ImageJ (NIH). For comparing cassette exon and retained intron splicing, band intensities were normalized to the adult isoform splice isoform to account for differences in EtBr intercalation for different length amplicons. Percent Spliced In (PSI) values were calculated for all alternative splicing events assayed per the following equation:

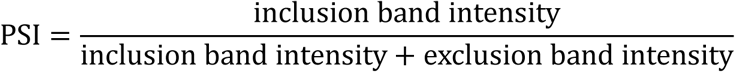

To determine relative steady-state and pre-mRNA abundance of *Clcn1*, probe-based quantitative RT-PCR (RT-qPCR) was used with primer/probe sets either pre-designed to target exon-exon borders for assaying *Clcn1* steady-state mRNA (IDT) or adapted to target intron-exon borders for assaying pre-mRNA levels. A pre-designed primer/probe set (IDT) for the TATA box binding protein (*Tbp*) was used as the reference/housekeeping gene for all reactions. All primer/probes are listed in Supplemental Table 1a with efficiencies greater than 90%. The Luna Universal Probe One-step RT-qPCR kit was used with 100ng total RNA input from each sample (isolation as described above) in 20uL reactions run with the Quantstudio 3 (Applied Biosystems, Thermo Fisher Scientific). The 2^−ΔΔ*CT*^ method was used to determine relative expression^59^. For all RT-PCR experiments, greater than 3 biological replicates were used for splicing analysis. All RT-qPCR experiments were completed with at least 3 biological replicates per genotype/timepoint in technical triplicate. Samples were standardized to the mean P18 WT cycle threshold (Ct) for samples on each plate.

### Long-read whole-Clcn1 sequencing library preparation

To analyze isoform utilization and splicing interactions at multiple locations in *Clcn1* transcripts we used long-read whole-transcript sequencing with the Pacific Biosciences Single Molecule Real Time (SMRT) system (Fig. S3a). This allowed for the generation of thousands of contiguous whole transcripts reads from exon 1 to exon 23 of *Clcn1* transcripts. Sequencing libraries were generated based on the PacBio protocol for multiplexing with barcoded universal primers. To start, RNA from quadriceps tissue from postnatal day 7 and 18 mice (3 biological replicates per genotype/timepoint) were reverse transcribed as discussed earlier. Two-microliters of each cDNA library was then used for target specific amplification (Phusion high-fidelity polymerase; NEB) of *Clcn1* transcripts with primers designed to bind within exon 1 and exon 23 with overhanging sequences and 5’AmMC6 blocking (Supplemental Table 1a). Samples run in parallel were amplified for 30 cycles and visualized on a 1% agarose gel (EtBr staining) for verification of amplicon identity and specificity. An annealing temperature of 63°C and 3% DMSO was used for first-step amplification. Samples were amplified for 20 cycles to progress in library preparation. These products were purified with AMPure XP beads (Beckman Coulter) at a volumetric ratio of 0.6 beads: 1 PCR product to select for the ∼3kb expected product. Next, 1.5uL of purified product was used for barcoding PCRs with Pacbio Universal Barcoding Primers (Pacific Biosciences). Barcoded products were then purified again with AMPure XP beads, quantified with Qubit high-sensitivity DNA assays (Invitrogen), and then equimolar pooled to enter the PacBio DNA repair and adapter ligation protocol (SMRTbell Express Template Prep Kit 2.0, Pacific Biosciences). The pooled library diluted and submitted for sequencing on the PacBio Sequel system at the Molecular Biology and Genomics Core (Washington State University, Pullman, WA).

### Long-read whole-*Clcn1* sequencing analysis

Initial processing of circular-consensus sequences (ccs), quality control, and demultiplexing was completed with the SMRT Link software package (Pacific Biosciences). Q20 reads were then processed with the pbBioconda (Pacific Biosciences) distribution using the lima package to remove overhangs and Isoseq to refine reads. Reads then entered the FLAIR analysis pipeline to align reads to chromosome 6 (mm39), define transcript isoforms present, and quantify isoform utilization^60^. The Refseq annotation of mm39 was used for determining known isoforms (UCSC distribution). Next, isoform utilization was quantified for each sample. A total of 275,185 barcoded Q20 reads were generated, with an average of 8,227 ± 4,709 reads per sample (mean±SD) with a median quality of Q38. To compare differential utilization of the various *Clcn1* isoforms in the different sample groups DESeq2 on R was utilized for isoforms with greater than 100 reads mapped across all samples^61^.

### Whole-cell voltage-clamp recordings of ClC-1 currents

To assay functional expression of ClC-1 channels in these mice, whole-cell patch-clamp experiments on isolated *flexor digitorum brevis* (FDB) muscle fibers were completed with solutions designed to differentiate ClC-1 chloride currents. Experiments were completed as previously outlined^17^. Briefly, mice were euthanized by cervical dislocation and decapitation, the FDB was excised from both hindlimbs, digested with 2mg/mL collagenase A (Roche) dissolved in standard Ringer’s solution for 45-60 minutes, and then dissociated by mechanical agitation with fire-polished pasture pipettes of decreasing bore diameter. Isolated fibers were plated on 35mm cell-culture dishes (Falcon) and stored at room temperature in Ringer’s solution until used within 8 hours of dissociation. Prior to experiments, Ringer’s solution was exchanged for external recording solution (in mM): 145 TEA-Cl, 10 CaCl_2_, 10 HEPES, 0.25 CdCl_2_, 0.003 nifedipine (pH 7.4 w/ TEA-OH). Then, whole-cell patch-clamp experiments were completed with low-resistance thin-walled pipettes (approximately 0.5 – 1.0 MOhms) were filled with (in mM): 130 Cs-aspartate, 10 CsCl, 5 MgCl_2_, 10 Cs_2_-EGTA, and 10 HEPES (pH 7.4 w/ CsOH). Solutions were designed to isolate Cl-currents (TEA/Cs to block K+ currents; no Na+ present; and nifedipine and Cd^2+^ to block Ca^2+^ currents) with a reversal potential for Cl-(VCl^−^) of approximately −53mV (Fig. 2-6). Recordings were collected with an Axopatch 200B (Molecular Devices) amplifier with voltage protocols and data acquisition controlled by pCLAMP 11 software (Molecular Devices). After formation of a gigaseal, whole-cell configuration was achieved, and fibers were held at −50mV for greater than five-minutes for equilibration. Series resistance was compensated (>90%), and currents were collected in response to voltage protocols featuring a pre-pulse to +60mV (250ms) for maximal activation of ClC-1 channels followed by a family of voltage-steps from −140 mV to +60 mV (Δ10mV; 500ms; 0.1Hz) to produce current-voltage (IV)-curves and tail currents for calculating relative open probability. To isolate ClC-1 currents, the external solution was then exchanged to also include 500μM 9-Anthracenecarboxylic acid (9-AC) to block ClC-1. Currents from the same protocol were then collected, representing all non-9-AC-sensitive currents (e.g., leak, capacitive, and non-ClC-1 Cl^−^ currents). Offline subtraction of traces collected before and after the addition of 9-AC yielded just 9-AC sensitive currents, corresponding to ClC-1 currents. To standardize across fiber sizes, a capacitive transient will be recorded from a 10mV depolarization from holding and integrated to calculate capacitance. Peak instantaneous and steady-state currents will then be divided by capacitance to yield current-densities (CD; pA/pF). Linear properties, including capacitance, access resistance, and membrane resistance were tracked with the Clampex software (Molecular Devices) throughout the collection of recordings. Preliminary analysis was completed with Clampfit software (Molecular Devices) with final analysis and plotting completed with Microsoft excel and GraphPad Prism 10 (Dotmatics). Representative traces were blanked to remove artifacts for presentation. Instantaneous currents were measured from voltage-step recordings immediately after the transition from holding at +60mV, whereas steady-state current was measured at the end of the 500ms voltage step. Instantaneous currents plotted versus voltage was fit with a modified Boltzmann equation for current at a specific membrane potential *I*(V_m_):

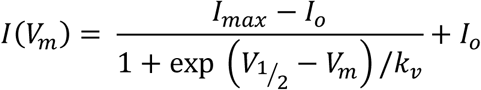

V_m_ is membrane potential, V_1/2_ voltage of half-maximal activation, k_v_ is slope factor, *I_max_* maximal current, and *I*_o_ is a constant offset current. Relative open probabilities based on normalized tail currents were fit to the equation of relative P_o_ at a given membrane potential: P_o,rel_(V_m_):

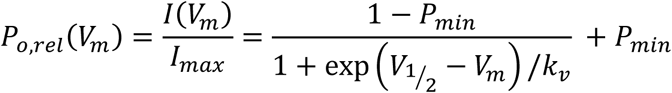

V_m_ is membrane potential, *I_max_* is the absolute maximal tail current, V_1/2_ voltage of half-maximal activation, P_min_ is the minimal value of P_o_ and k_v_ is slope factor.

### Rotarod

To assay the presence of myotonia and general muscle function rotarod experiments with a linear acceleration protocol were completed, as previously described^19^. Briefly, on day-one of experiments mice (>5 mice/genotype) ten-to-twelve weeks-postnatal were first acclimated to the rotarod (Columbus Instruments) spinning at 4 rotations-per-minute (rpm) for three 1-minute trials separated by at least fifteen minutes. Mice were then subjected to four-trials of linear-acceleration protocols starting at 4rpm and accelerating to 40rpm over 5-minutes at which time the rotarod continues to spin at a constant 40rpm for an additional 5-minutes. Time-to-fall (seconds) and rotation speed (rpm) at which the mouse fell was recorded for each trial. At least 15-minutes of rest was provided between trials. The mice were then trained for another two-days with 4-trials of linear acceleration protocol (3-days training total). Formal data collection was completed on the fourth day with 4-trials of linear acceleration protocol. The average of the 4-trials for each mouse was used for analysis.

### *In vivo* muscle-contraction with paired electromyography (EMG)

To assay in vivo force production, myotonia, and underlying electrical activity, a standard *in vivo* force contraction assay of murine plantarflexion was used with the addition of EMG recordings^62^. Mice (approximately 4-months postnatal; >5 mice/genotype) were aestheticized with inhaled isoflurane, hair on the hind-limb assayed was shaved, and the mouse was placed on a heated (37°C) platform with the knee fixed to immobile posts. Next, the footpad was securely attached to a footplate connected to a force-transducer to measure plantarflexion force (Aurora). Two-stimulating electrodes were then placed on the medial aspect of the upper-hindlimb to stimulate the tibial nerve and produce contraction of the posterior compartment of the lower hindlimb (primarily the gastrocnemius). Stimulating electrode placement and current amplitude was optimized to produce the peak force in response to single 1.25 millisecond (ms) test pulses. Next, a protocol run delivered that featured 3 twitch contractions (0.2ms) and a single tetanus (150Hz for 500ms). Following the acquisition of these recordings, monopolar EMG electrodes (iWorx Systems) were placed subcutaneously as close to the gastrocnemius muscle belly as possible at proximal and distal locations for the collection EMG data; electrical activity in response to the same contraction protocol was collected secondary to the initial force recordings, as the presence of additional EMG electrodes suspended in the distal hindlimb provided artifacts in the force recordings. Additionally, a ground electrode was placed subcutaneously in the abdominal region. Data was collected and amplified with an IX-BIO 4 isolated biopotential recorder then acquired with an IX-RA-5x attached to a workstation running the LabScribe software for data collection (iWorx Systems). The same protocol of three twitches followed by a single tetanus was delivered by the tibial nerve stimulating electrodes and electrical activity was recorded with the LabScribe software (iWorx). At least 5 minutes rest was provided between tetanus stimulations to prevent warm-up. Force recordings from the initial tetanus was measured for peak force calculations (normalized to mouse bodyweight) and myotonia (impulse of normalized force trace; time integral of normalized force). EMG signals were aligned based on the initial takeoff of the voltage signal in response to the stimulus artifact. For summarizing the overall electrical activity, of the muscle a linear envelope was calculated by rectifying and applying a second-order low-pass Butterworth filter with MatLab (MathWorks^63^). Fast Fourier Transform was utilized to evaluate amplitude/power spectra.

### *Ex vivo* / *In vitro* muscle-contraction

Force measurements were collected from the soleus excised from 4-6-month postnatal mice (>5 mice/genotype). Established protocols were used for assaying muscle isometric force generation, frequency dependence, and myotonia, as previously described^10,64^. First, mice were anesthetized with inhaled isoflurane (∼2%). *Soleus* muscles were carefully dissected and excised with suture loops tied to both proximal and distal tendons for later attachment to the immobile post and force transducer, respectively. Muscles were then submerged in a bath between two platinum stimulating electrodes in oxygenated (95% O_2_ and 5% CO_2_) and warmed Ringer’s solution. Ringer’s solution contained (in mM): 1.2 NaH_2_PO_4_, 1 MgSO_4_, 4.83 KCl, 137 NaCl, 2 CaCl_2_, 10 glucose, 24 NaHCO_3_ at pH 7.4. Force was measured with an Aurora Scientific 1200A *ex vivo* system equipped with an 809B muscle testing system with a 300C-LR force transducer and a 701C stimulator (Aurora Scientific). Muscles were equilibrated in the bath for at least 10 minutes prior to recordings. Optimal length (L_o_) of the muscle and stimulation current was determined for each muscle. The first a stimulation protocol was the applied for determination of twitch and tetanic contractile properties (peak force and myotonia); three twitches (0.2ms simulation; 20s between stimulations) and three tetani (500ms of 0.2ms stimuli delivered at 150Hz) were recorded. Next, frequency dependence of force production was assayed with a standard protocol of increasing stimulation frequencies, as appropriate for each muscle group. All force recordings reported were normalized to the cross-sectional area (CSA) through measurement of muscle length and weight as well as the use of the appropriate pyknotic vector for each muscle and thus reported as specific force (mN/mm^2^). For frequency dependence, force was normalized to the peak force generated by the muscle across all the stimuli; therefore, reported as normalized force (arbitrary unit, AU). Similarly, for assaying myotonia through impulse calculation, peak force generated during the tetanic stimuli was normalized to 1 AU. The time integral of the force trace following the tetanic stimuli was then calculated to provide the area under curve (impulse) of myotonia.

### Histology – conventional staining and immunohistochemistry

For analysis of skeletal muscle histopathologic hallmarks of DM1, standard hematoxylin and eosin (H&E) was used for assessing central nucleation. Muscles were mounted with tragacanth gum, frozen in 2-methylbutane submerged in liquid nitrogen, and stored at - 80°C until further processing. Transverse slices (10um) were obtained with a cryostat (Thermo Scientific) and transferred to slides for staining such that each slide contained at least one-slice of each genotype assayed. Slides were stored at −80°C prior to staining. For determination of fiber-type composition, myosin heavy chain (MyHC) immunohistochemistry (IHC) was used for visualization of type I, IIa, IIb, and IIx fibers. Slices were first permeabilized for 10-minutes with PBST (PBS with 0.2% Triton X-100) and then blocked at room temperature for 30-minutes in 10% normal goat serum (NGS) (Gibco, Thermo Fisher Scientific) in PBS. Additional blocking with 0.1mg/mL AffiniPure Fab fragment anti–mouse IgG (3%; H&L) (Jackson Immunoresearch) in PBS with 2% normal goat serum for 60-minutes (room temperature). Slides were then washed (3 times with fresh PBS for 10-mins at room temperature) and incubated with primary antibodies (Supplemental Table 1b) overnight (14-18 hours) at 4°C; primary antibodies against myosin heavy chain isoforms were obtained from the Developmental Studies Hybridoma Bank (DSHB) created by the NICHD of the NIH and maintained at The University of Iowa, Department of Biology (Iowa City, IA). The next day, slides were washed again with PBS (3 times for 10-mins at room temperature). Then, secondary antibodies were applied for one-hour at room temperature. Slides were then washed with PBS (3 times for 10-mins at room temperature) and finally mounted with Fluoromount-G (Invitrogen). No primary antibody control slides were completed for all samples with all IHC staining paradigms and were used for establishing exposure levels when imaging each slide. Imaging was performed with a Keyence BZ-X810 All-in-One Florescence Microscope. Magnification and scale are noted for all included images. Percent fibers with central nuclei were manually calculated as the number of fibers with 1+ centrally located nuclei divided by the total number of intact fibers in a complete H&E-stained transverse section (image collected at 10x and stitched with Keyence analysis software). Individuals performing quantification were blinded to genotype when possible. Fiber morphometry and typing was performed with Myosoft^65^. For presentation of minimal Feret’s diameter and CSA, values for each muscle sample were filtered for outliers to remove rare errors of the automated analysis software by the ROUT method (Q = 1%) with Graphpad Prism 10. For presentation of representative images, ImageJ (NIH) with the plugin *Quick Figures* was utilized^66^.

### RNA sequencing – data collection, preliminary data processing, and expression analysis

RNA was isolated from quadriceps muscle from 4–6-month-old WT, ClC-1^ΔE7a/ΔE7a,^ Mbnl1^−/−^, and Mbnl1^−/−^ ; ClC-1^ΔE7a/ΔE7a^ mice as described above (RNA isolation). Raw reads generated from the Illumina base calls were demultiplexed using bcl2fastq version 2.20.0. Quality filtering and adapter removal are performed using FastP version 0.23.1 with the following parameters: “--length_required 35 --cut_front_window_size 1 -- cut_front_mean_quality 13 --cut_front --cut_tail_window_size 1 --cut_tail_mean_quality 13 --cut_tail -y –r“^67^. Processed/cleaned reads were then mapped to the GRCm39/gencode M31 reference using STAR_2.7.9a with the following parameters: “— twopass Mode Basic --runMode alignReads --outSAMtype BAM Unsorted – outSAMstrandField intronMotif --outFilterIntronMotifs RemoveNoncanonical – outReadsUnmapped Fastx“^68,69^. Gene level read quantification was derived using the subread-2.0.1 package (featureCounts) with a GTF annotation file (GRCm39/gencode M31) and the following parameters for stranded RNA libraries “-s 2 -t exon -g gene_name“^70^. Differential expression analysis was performed using DESeq2^61^ with a minimum threshold of 50 aggregate reads per gene. PCA analysis was completed with pcaExplorer^71^. Heatmaps were generated with ComplexHeatmap^72^. Venn diagrams were created with Bioinformatics and Evolutionary Genomics web tool (https://bioinformatics.psb.ugent.be/webtools/Venn/).

### Short-read mRNA-sequencing splicing analysis

Isolated RNA was sequenced to a depth of ∼60 million reads and mapped against the GRCm39 mouse genome using the STAR 2.7.0 aligner. The aligned reads were analyzed for splicing alterations using MAJIQ and ΔPSI-values were calculated in pair-wise comparisons for all conditions with of ΔPSI > 0.1 or < −0.1 and an FDR < 0.05 were set to be significantly changed. For visualization, MAJIQ modulizer results separated after event types (exon skipping, intron retention, alternative 3’ splice sites and 5’splice sites) were loaded into R 4.2. Partial rescues were defined as having more than a 0.15 delta-PSI shift in Mbnl1^−/−^; ClC-1^ΔE7a/ΔE7a^ towards control as compared to Mbnl1^−/−^ mice. Events were considered to display a complete rescue with ΔPSI > 0.1 or < −0.1 in Mbnl1^−/−^ vs WT mice and ΔPSI < 0.1 or > −0.1 (non-changing) in Mbnl1^−/−^ ; ClC-1^ΔE7a/ΔE7a^ vs WT at an FDR of < 0.05).

### Statistics

Statistical analysis and graphing were predominately completed with GraphPad Prism 10. For RNA-seq and long-read PacBio sequencing, statistics with DESeq2 were completed with R (4.3.1). Some illustrations and schematics were generated with BioRender (Biorender.com). For comparisons of more than two groups with parametric data that features reasonable sphericity, standard one-way ANOVAs with secondary multiple comparisons with Tukey’s correction were completed. For comparisons with clear differences in variance between groups, Brown-Forsythe and Welch ANOVA tests with secondary multiple comparisons with Dunnett T3 correction was completed. For non-parametric comparisons, a Kruskal-Wallis test with Dunn’s multiple comparisons were completed. Comparison between genotypes across different timepoints was completed with two-way ANOVA with Tukey’s corrected multiple comparisons. Statistics for long-read sequencing and RNAseq analysis were completed as discussed in respective methods sections. For all comparisons shown, significance is indicated as follows unless otherwise specified: multiplicity adjusted p-value <0.05*, <0.01**, <0.001***, and <0.0001****.

## Resource availability

Requests for further information and resources should be directed to and will be fulfilled by the lead contact, John Lueck

## Materials availability

Materials will be made available upon request.

## Data and code availability

All data are presented in the manuscript and the study does not report original code. Any additional information required to reanalyze the data reported in this paper is available from the lead contact upon request.

## Acknowledgments

We thank members of the Lueck laboratory for reading and commenting on the manuscript. We gratefully acknowledge the Genome Editing Core at Augusta University for generating the ClC-1^Δe7a^ mouse model necessary to conduct this research. We wish to thank Norma Sinclair, Rongbin Guan, and Joanne Schwarting for their technical expertise in generating transgenic mice. We also wish to thank Jacqueline Heller, the Klein Family and the Marigold Foundation for their generous support. This study was funded by NIH grants 1F31AR082284 (to M.T.S.) and T90DE021985 and T32GM135134 (to LAC); Doctoral Fellowships from The Myotonic Dystrophy Foundation (to L.A.C. and S.H.); and NIH grant R01AR079424 (to JDL). P.M. is supported by the Deutsche Forschungsgemeinschaft (DFG, German Research Foundation – 470092532).

