## Supplemental Figures for "Elimination of myotonia improves myopathy in a muscleblind knockout model of myotonic dystrophy"

### Supplemental Figures and Tables

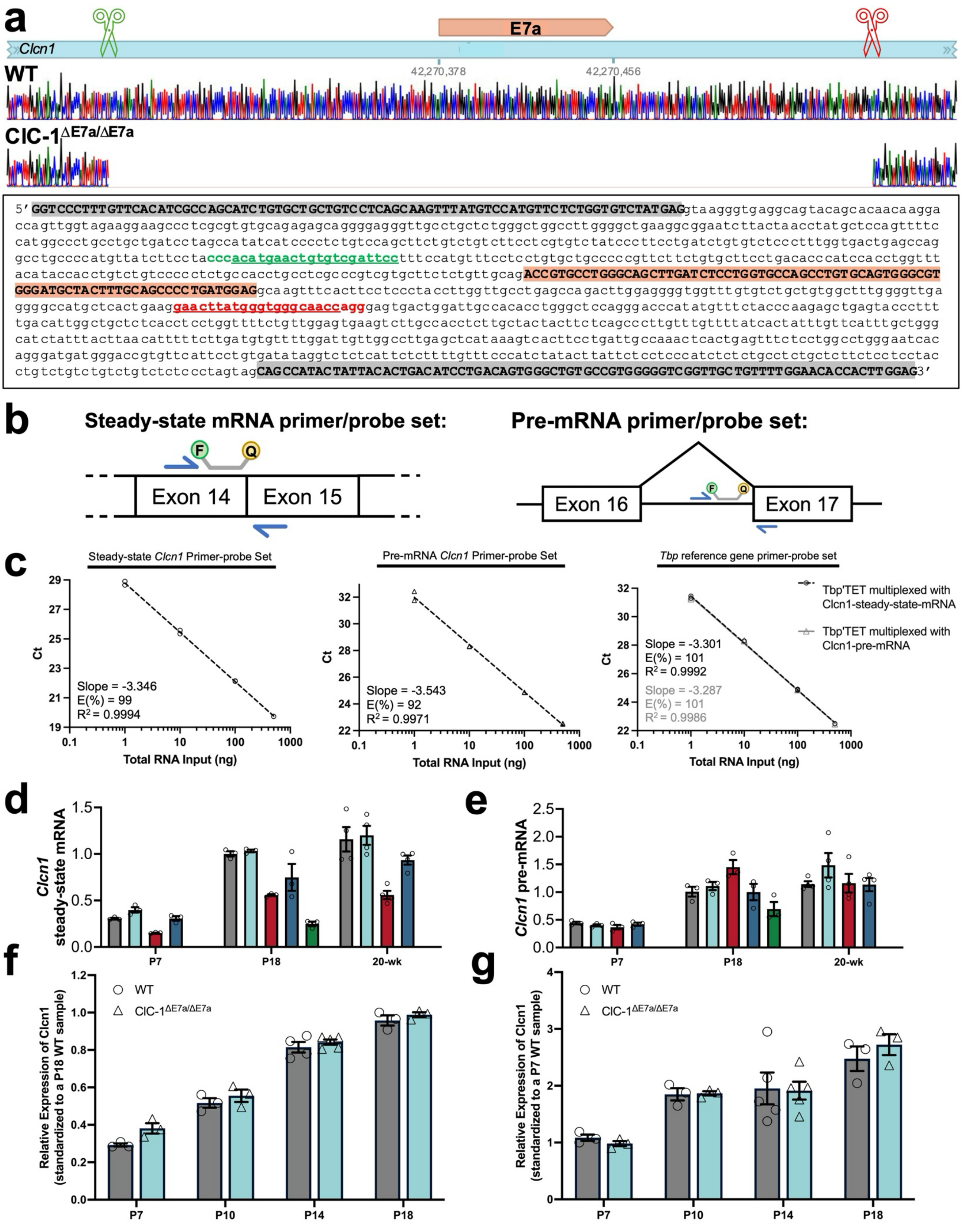
a
E7a
Clcn1
42,270,378
42,270,456
WT
ClC-1ΔE7a/ΔE7a
5′ GGTCCCTTTGTTCACATCGCCAGCATCTGTGCTGCTGTCCTCAGCAAGTTTATGTCCATGTTCTCTGGTGTCTATGAGgtaagggtgaggcagtacagcacaacaagga
ccagttggtagaaggaagccctcgcgtgtgcagagagcaggggagggttgcctgctctgggctggccttggggctgaaggcggaatcttactaacctatgctccagttttc
catggccctgcctgctgatcctagccatatcatcccctctgtccagcttctgtctgtcttcctcgtgtctatcccttcctgatctgtgtctccctttggtgactgagccag
gcctgccccatgttatcttcctacccacatgaactgtgtcgattcctttccatgtttcctcctgtgctgccccgttcttctgtgcttcctgacacccatccacctggttt
acataccacctgtctgtcccctctgccacctgcctcgcccgtcgtgcttctctgttgcagACCGTGCCTGGGCAGCTTGATCTCCTGGTGCCAGCCTGTGCAGTGGGCGT
GGGATGCTACTTTGCAGCCCCTGATGGAGgcaagtttcacttcctccctaccttggttgcctgagccagacttggaggggtggtttgtgtctgctgtggctttgggttga
ggggccatgctcactgaaggaacttatgggtgggcaaccagggagtgactggattgccacacctgggctccagggacccatatgtttctacccaagagctgagtaccctttt
tgacattggctgctctcacctcctggtttctgttggagtgaagtcttgccacctcttgctactacttctcagcccttgtttgttttatcactatttgttcatttgctggg
catctatttacttaacatttttcttgatgtgttttggattgttggccttgagctcataaagtcacttcctgattgccaaactcactgagtttctcctggcctgggaatcac
agggatgatgggaccgtgttcattcctgtgatataggtctctcattctctttgtttcccatctatacttattctcctcccatctctctgcctctgctcttctcctcctac
ctgtctgtctgtctgtctctccctagtagCAGCCATACTATTACACTGACATCCTGACAGTGGGCTGTGCCGTGGGGGTCGGTTGCTGTTTTGGAACACCACTTGGAG3′
b
Steady-state mRNA primer/probe set:
F
Q
Exon 14
Exon 15
Pre-mRNA primer/probe set:
Exon 16
Exon 17
c
Steady-state Clcn1 Primer-probe Set
Ct
Slope = -3.346
E(%) = 99
R2 = 0.9994
Total RNA Input (ng)
Pre-mRNA Clcn1 Primer-probe Set
Slope = -3.543
E(%) = 92
R2 = 0.9971
Tbp reference gene primer-probe set
Tbp'TET multiplexed with Clcn1-steady-state-mRNA
Tbp'TET multiplexed with Clcn1-pre-mRNA
Slope = -3.301
E(%) = 101
R2 = 0.9992
Slope = -3.287
E(%) = 101
R2 = 0.9986
0.1
1
10
100
1000
d
Clcn1 steady-state mRNA
P7
P18
20-wk
e
Clcn1 pre-mRNA
f
WT
ClC-1ΔE7a/ΔE7a
Relative Expression of Clcn1 (standardized to a P18 WT sample)
P7
P10
P14
P18
g
Relative Expression of Clcn1 (standardized to a P7 WT sample)

SF 1

**Figure S1** Generation of Clcn1 exon 7a exclusion mouse and RT-qPCR assessment of *Clcn1.* (a) Verification of genomic excision of exon 7a sequence (and flanking intronic regions) from *Clcn1.* Upper panel shows electropherogram of sanger sequencing of targeted amplification of the *Clcn1* gene from genomic DNA isolated from WT and ClC-1$^{\Delta E7a/\Delta E7a}$ samples, aligned to the *Clcn1* sequence above with CRISPR cut sites indicated by scissors and the exon 7a sequence annotated. Bottom panel represents the *Clcn1* genomic sequence flanking exon 7a (highlighted in red) with exons 6 and 7 shaded in gray and CRISPR cut sequences in green and red text. Electropherogram and schematic generated with Benchling. (b) Schematics of the design of RT-qPCR primer-probe sets for assessing steady-state mature *Clcn1* mRNA (IDT) and *Clcn1* pre-mRNA. (c) Standard curves to assess primer probe efficiencies estimated by dilution series on total RNA isolated from P14 wildtype mouse (completed in technical triplicate). (d) Relative steady-state *Clcn1* and (e) *Clcn1* pre-mRNA used for calculation of steady-state to pre-mRNA ratio in Fig. 1d. (f) Relative steady-state *Clcn1* and (g) *Clcn1* pre-mRNA across early post-natal development. Tbp as housekeeping. Normalized to P18 WT average or P7 WT average. Dots represent biological replicates. Completed in technical triplicate. Data shown mean ± SE.

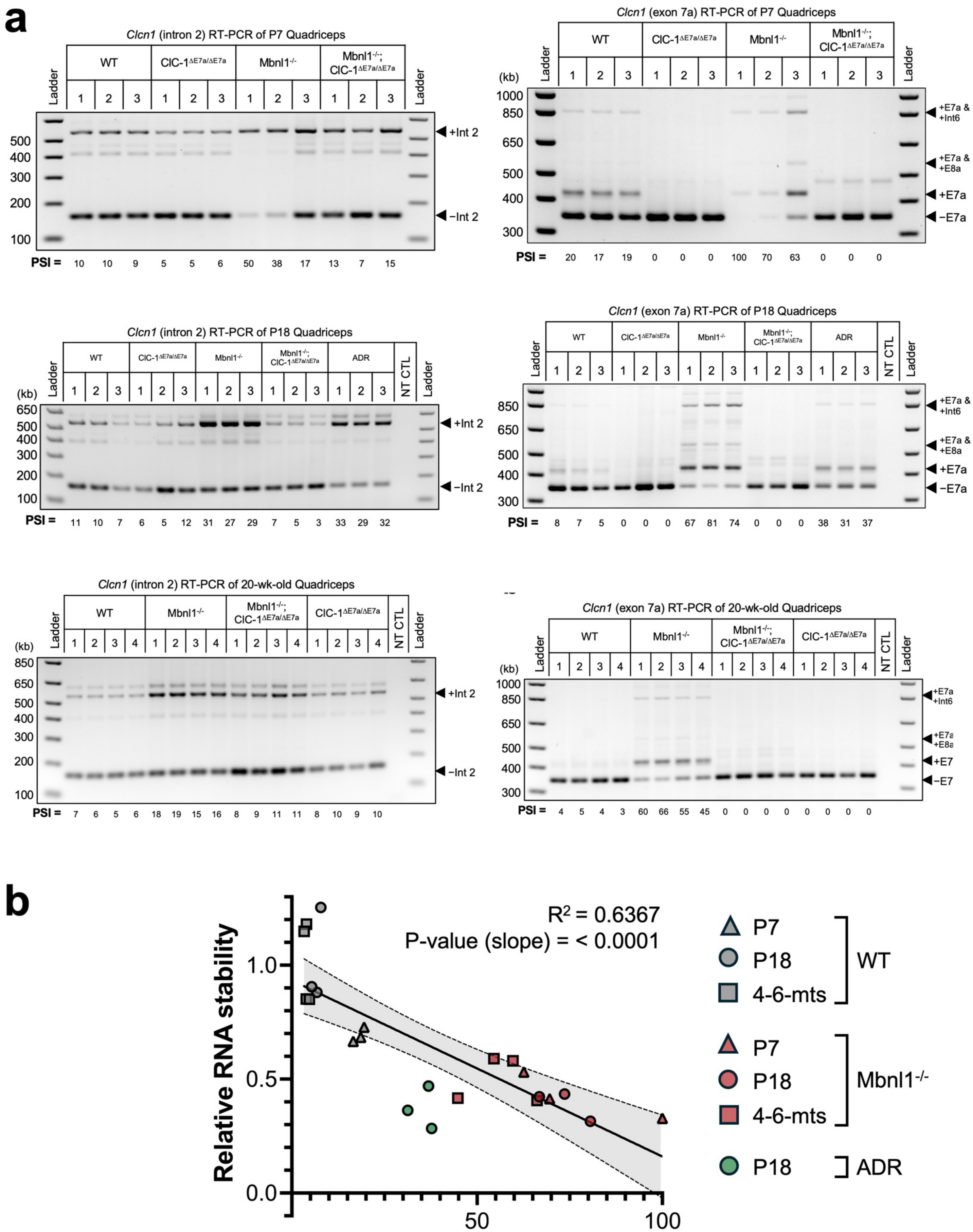
a
Clcn1 (intron 2) RT-PCR of P7 Quadriceps
Ladder
WT
ClC-1ΔE7a/ΔE7a
Mbnl1-/-
Mbnl1-/-; ClC-1ΔE7a/ΔE7a
500
400
300
200
100
+Int 2
−Int 2
PSI = 10 10 9 5 5 6 50 38 17 13 7 15
Clcn1 (exon 7a) RT-PCR of P7 Quadriceps
(kb)
1000
850
650
500
400
300
+E7a & +Int6
+E7a & +E8a
+E7a
−E7a
PSI = 20 17 19 0 0 0 100 70 63 0 0 0
Clcn1 (intron 2) RT-PCR of P18 Quadriceps
ADR
NT CTL
650
PSI = 11 10 7 6 5 12 31 27 29 7 5 3 33 29 32
Clcn1 (exon 7a) RT-PCR of P18 Quadriceps
PSI = 8 7 5 0 0 0 67 81 74 0 0 0 38 31 37
Clcn1 (intron 2) RT-PCR of 20-wk-old Quadriceps
850
PSI = 7 6 5 6 18 19 15 16 8 9 11 11 8 10 9 10
Clcn1 (exon 7a) RT-PCR of 20-wk-old Quadriceps
PSI = 4 5 4 3 60 66 55 45 0 0 0 0 0 0 0 0
b
Relative RNA stability
Exon 7a PSI (%)
0.0
0.5
1.0
50
100
R2 = 0.6367
P-value (slope) = < 0.0001
P7
P18
4-6-mts
WT
Mbnl1-/-
ADR

SF2

**Figure S2** Representative RT-PCR gels and relationship between exon 7a splicing and relative RNA stability. (a) Representative agarose gels stained with EtBr for targeted RT-PCR splicing analysis of intron 2 retention and exon 7a inclusion for *Clcn1* with WT, ClC-$1^{\Delta E7a/\Delta E7a}$, Mbnl1$^{-/-}$, Mbnl1$^{-/-}$ ; ClC-$1^{\Delta E7a/\Delta E7a}$, and ADR samples as indicated at P7, P18, and 4-6-month time-points. Lanes represent biological replicates. Ladder (1kb Plus ladder, Invitrogen) and no template control (NT CTL) shown. Gel for P18 exon 7a splicing is reproduced from Fig. 1b for completeness. (b) Linear regression analysis of samples plotted with exon 7a PSI against relative RNA stability, for all samples without forced exon 7a exclusion. Line represents simple linear regression with shaded area representing 95% CI.

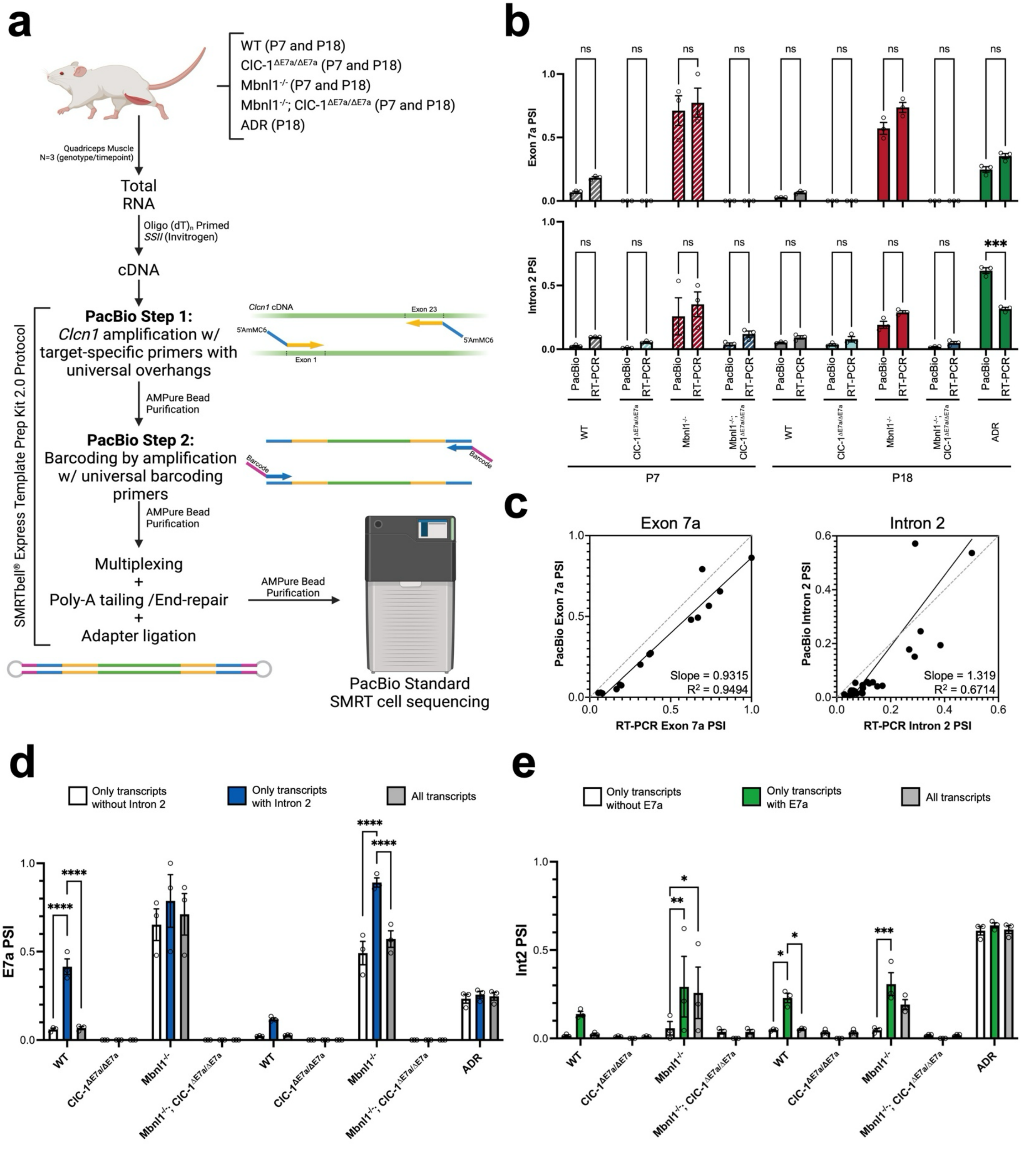
a
WT (P7 and P18)
ClC-1ΔE7a/ΔE7a (P7 and P18)
Mbnl1-/- (P7 and P18)
Mbnl1-/-; ClC-1ΔE7a/ΔE7a (P7 and P18)
ADR (P18)
Quadriceps Muscle N=3 (genotype/timepoint)
Total RNA
Oligo (dT)n Primed SSII (Invitrogen)
cDNA
SMRTbell® Express Template Prep Kit 2.0 Protocol
PacBio Step 1: Clcn1 amplification w/ target-specific primers with universal overhangs
Clcn1 cDNA
Exon 23
Exon 1
5'AmMC6
AMPure Bead Purification
PacBio Step 2: Barcoding by amplification w/ universal barcoding primers
Barcode
AMPure Bead Purification
Multiplexing + Poly-A tailing /End-repair + Adapter ligation
AMPure Bead Purification
PacBio Standard SMRT cell sequencing
b
Exon 7a PSI
Intron 2 PSI
ns
***
PacBio
RT-PCR
WT
ClC-1ΔE7a/ΔE7a
Mbnl1-/-
Mbnl1-/-; ClC-1ΔE7a/ΔE7a
ADR
P7
P18
c
Exon 7a
PacBio Exon 7a PSI
RT-PCR Exon 7a PSI
Slope = 0.9315
R² = 0.9494
Intron 2
PacBio Intron 2 PSI
RT-PCR Intron 2 PSI
Slope = 1.319
R² = 0.6714
d
Only transcripts without Intron 2
Only transcripts with Intron 2
All transcripts
E7a PSI
****
WT
ClC-1ΔE7a/ΔE7a
Mbnl1-/-
Mbnl1-/-; ClC-1ΔE7a/ΔE7a
ADR
e
Only transcripts without E7a
Only transcripts with E7a
All transcripts
Int2 PSI
*
**
***

**SF 3**

**Figure S3** Long-read *Clcn1* sequencing and comparison to traditional RT-PCR splicing analysis. (a) Schematic of targeted whole-*Clcn1* transcript PacBio sequencing. (b) Comparison between exon 7a and intron 2 PSI values derived from PacBio transcript utilization analysis and traditional RT-PCR with band densitometry. Comparisons represent two-sample t-tests with ”ns” representing $p \geq 0.05$ and *** $p < 0.001$. (c) Linear regression analysis of samples plotted against PacBio and RT-PCR determined PSI values for exon 7a inclusion (left) and intron 2 retention (right). Simple linear regression shown by solid black line with slope and R-squared values as indicated. Dotted line represents a slope of 1 (y=x). (d) Combinatorial splicing analysis of full-length *Clcn1* transcripts displaying exon 7a PSI based on intron 2 status of the transcripts (only transcripts without intron 2, only transcripts with intron 2, or all transcripts). (e) Combinatorial splicing analysis of full-length *Clcn1* transcripts displaying intron 2 PSI based on exon 7a status of the transcripts (only transcripts without exon 7a, only transcripts with exon 7a, or all transcripts). Presented mean ± SE with superimposed biological replicates at dots. Analysis completed with two-way ANOVA (genotype and splicing status) with intra-genotype comparisons between splicing status shown. Multiplicity adjusted p-value shown as * $p < 0.05$, ** $p < 0.01$, *** $p < 0.001$, **** $p < 0.0001$.

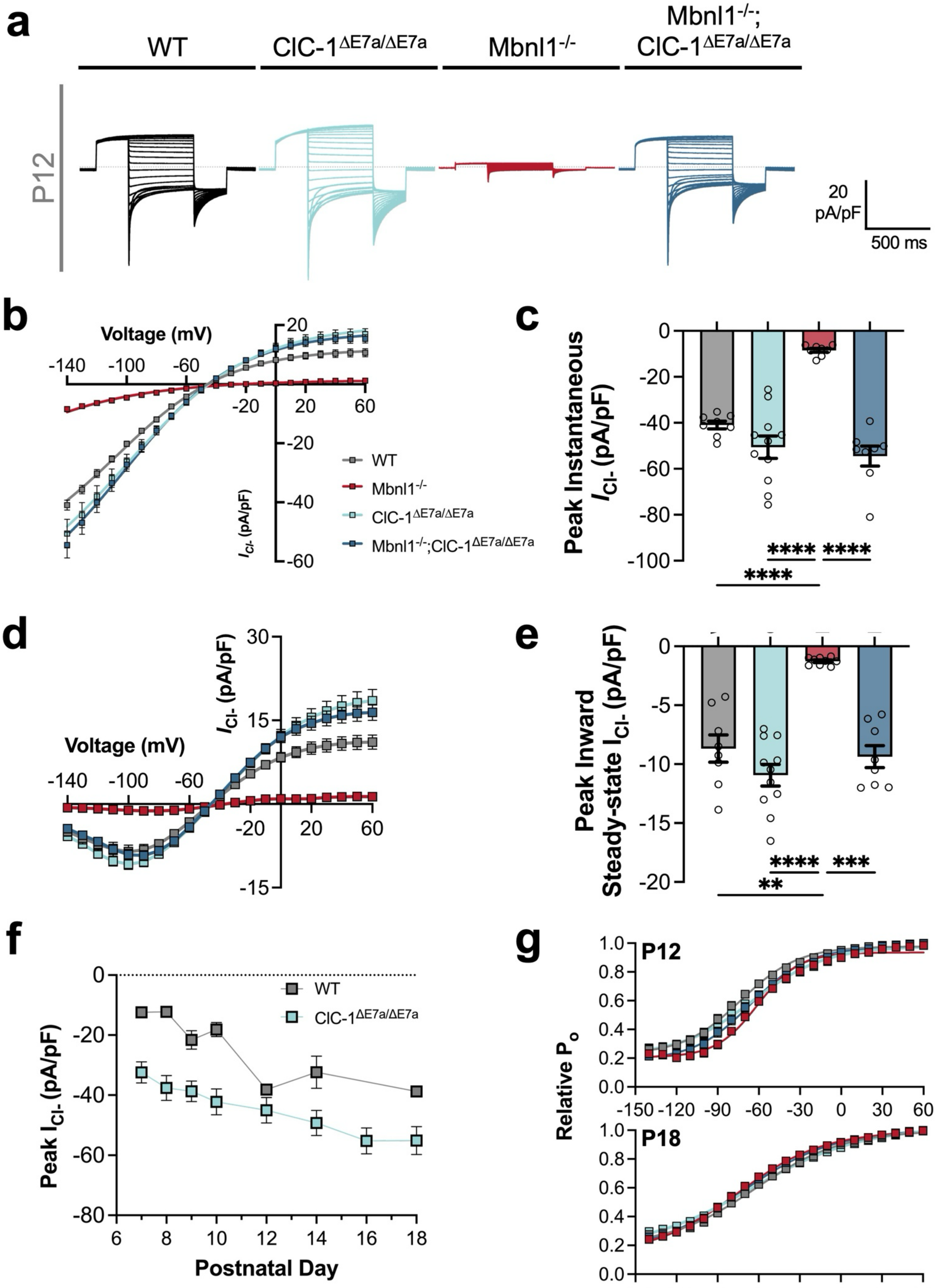
a
WT
ClC-1$^{ΔE7a/ΔE7a}$
Mbnl1$^{-/-}$
Mbnl1$^{-/-}$; ClC-1$^{ΔE7a/ΔE7a}$
P12
20 pA/pF
500 ms
b
Voltage (mV)
$I_{Cl-}$ (pA/pF)
WT
Mbnl1$^{-/-}$
ClC-1$^{ΔE7a/ΔE7a}$
Mbnl1$^{-/-}$;ClC-1$^{ΔE7a/ΔE7a}$
c
Peak Instantaneous $I_{Cl-}$ (pA/pF)
****
****
****
d
Voltage (mV)
$I_{Cl-}$ (pA/pF)
e
Peak Inward Steady-state $I_{Cl-}$ (pA/pF)
****
***
**
f
Peak $I_{Cl-}$ (pA/pF)
WT
ClC-1$^{ΔE7a/ΔE7a}$
Postnatal Day
g
P12
P18
Relative $P_o$
Voltage (mV)

SF 4

**Figure S4** Whole-cell patch-clamp of ClC-1 currents. (a) Representative traces of ClC-1 currents from WT, ClC-1$^{\Delta E7a/\Delta E7a}$, Mbnl1$^{-/-}$, and Mbnl1$^{-/-}$ ; ClC-1$^{\Delta E7a/\Delta E7a}$ samples collected at P12. Scale as shown. (b) Current density versus voltage for peak instantaneous inward current in response to family of voltage steps (-140 to 60 mV) with peak inward current for each P12 recording compared in (c) as a bar graph. (d) Average steady-state current versus voltage with max inward steady-state current compared in (e) as a bar graph. Comparisons with one-way ANOVA with post-hoc multiple comparisons. Multiplicity adjusted p-values shown as ** $p < 0.01$, *** $p < 0.001$, and **** $p < 0.0001$. (f) ClC-1 current densities across early post-natal development (P7 to P18) for WT and ClC-1$^{\Delta E7a/\Delta E7a}$ samples (n= 2+ mice with multiple fibers per mouse per timepoint per genotype). (e) Relative open probability versus voltage as determined by amplitude of tail currents for P12 and P18 samples fit with modified Boltzmann equation. Data shown mean ± SE. Linear properties and values of Boltzmann fit shown in supplemental table 2.

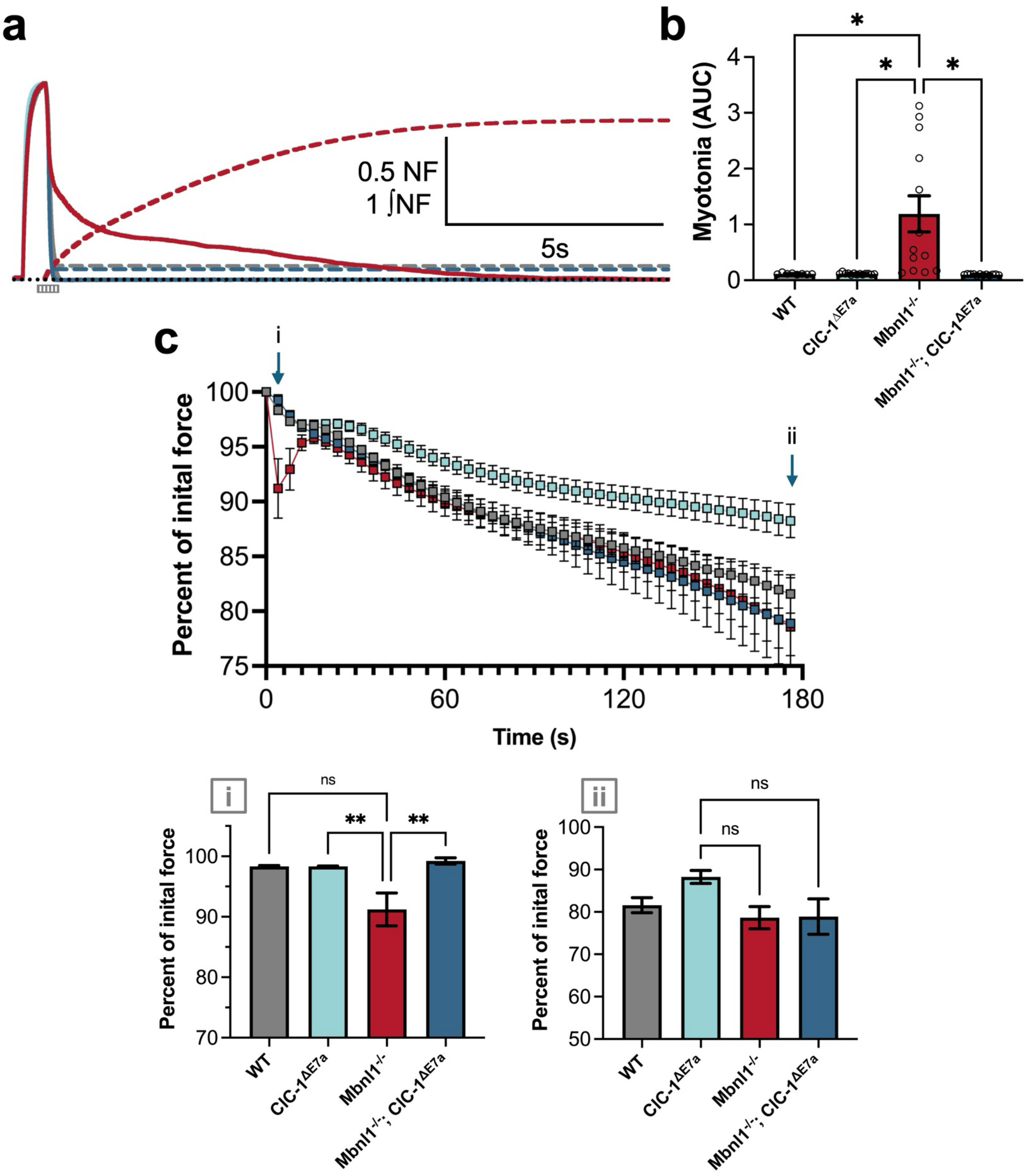
a
0.5 NF
1 ∫NF
5s
b
Myotonia (AUC)
WT
ClC-1ΔE7a
Mbnl1-/-
Mbnl1-/-; ClC-1ΔE7a
c
Percent of inital force
Time (s)
i
ii
ns
**
Percent of inital force
WT
ClC-1ΔE7a
Mbnl1-/-
Mbnl1-/-; ClC-1ΔE7a

SF 5

**Figure S5** *Ex vivo* force-contraction from isolated solei muscles. (a) Representative normalized force traces in response to tetanic stimuli (500 ms, 150Hz) with area under the myotonic force represented by dotted lines. Scales as shown. (b) Myotonia (AUC) compared for samples (arbitrary units). Comparison with one-way ANOVA with Tukey's multiple comparisons. Multiplicity adjusted p-values shown as * $p < 0.05$. (c) Peak force per tetanic stimuli for transient weakness and fatigue protocol comprised of 45 consecutive tetani (500 ms, 100 Hz) delivered every 4 seconds (duty cycle = 4s). Colors for each genotype as per bar graph boxes. Comparisons between groups for the second stimuli (i) and last stimuli (ii) to evaluate for transient weakness and fatigue, respectively, shown below. Analyzed with two-way ANOVA (stimulation number and genotype) with multiple comparisons with multiplicity adjusted p-values shown as ** $p < 0.01$. Data presented as mean ± SE.

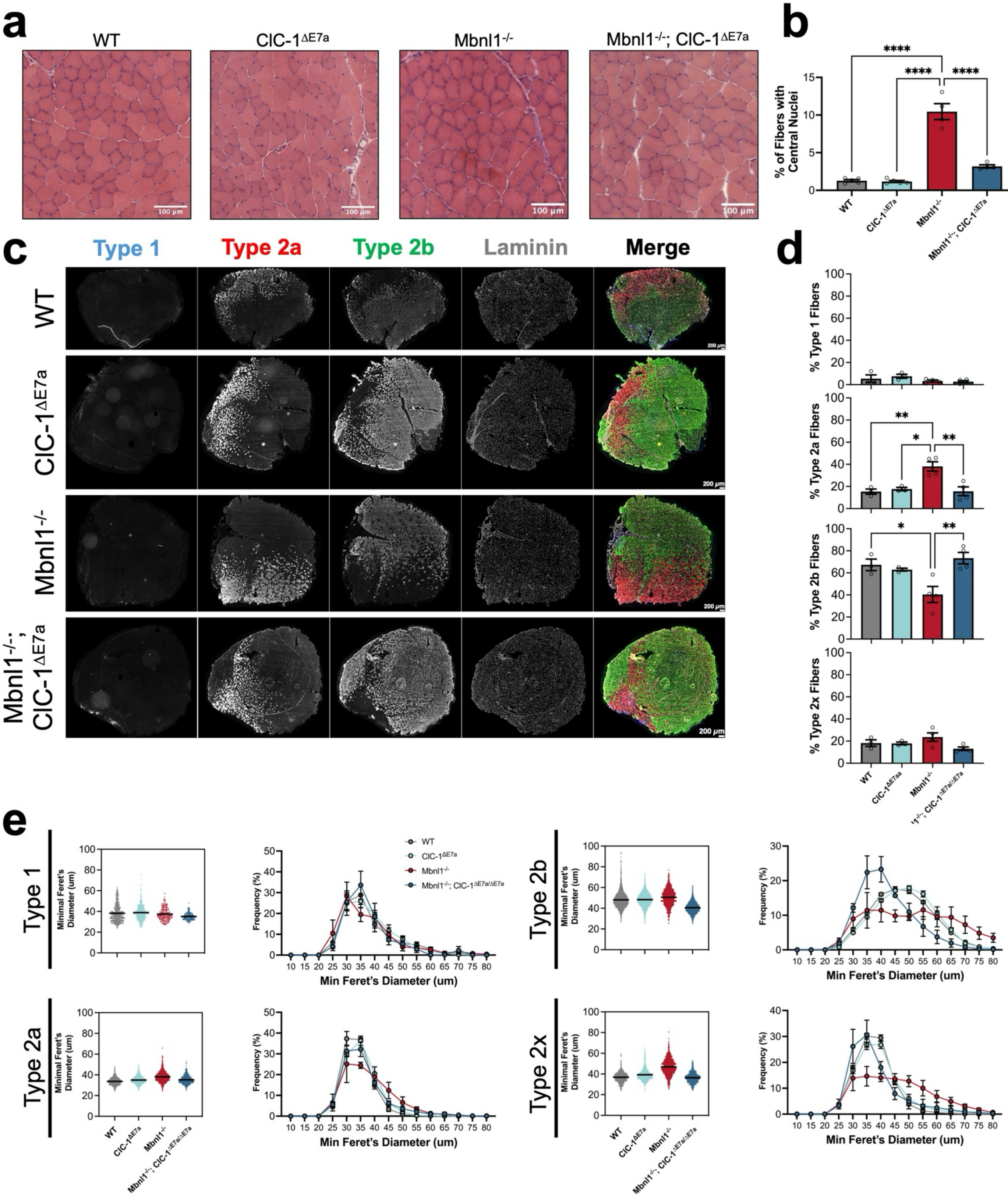
a
WT
ClC-1ΔE7a
Mbnl1-/-
Mbnl1-/-; ClC-1ΔE7a
100 µm
b
% of Fibers with Central Nuclei
****
c
Type 1
Type 2a
Type 2b
Laminin
Merge
200 µm
d
% Type 1 Fibers
% Type 2a Fibers
% Type 2b Fibers
% Type 2x Fibers
e
Minimal Feret's Diameter (um)
Frequency (%)
Min Feret's Diameter (um)
Mbnl1-/-; ClC-1ΔE7a/ΔE7a

SF 6

**Figure S6** Histological analysis of quadriceps muscle. (a) Representative images of quadriceps transverse sections stained with hematoxylin and eosin (H&E) from WT, WT, ClC-1$^{\Delta E7a/\Delta E7a}$, Mbnl1$^{-/-}$, and Mbnl1$^{-/-}$ ; ClC-1$^{\Delta E7a/\Delta E7a}$ mice (4-6-month-old). Scale bars shown represent 100 micrometers. (b) Percent of fibers with central nuclei present on H&E strained quadriceps samples. (c) Fiber-typing by myosin heavy chain (MyHC) isoform was completed by immunohistochemistry, representative images shown, with each MyHC isoform channel and laminin counterstain shown in grayscale and with the merged image pseudo-colored (based on color of channel labels). (d) Percent of fibers staining for each of the respective MyHC isoforms. Comparisons with one-way ANOVAs with Tukey's multiple comparisons. Multiplicity adjusted p-values of significant comparisons presented as * p<0.05, ** p<0.01, *** p<0.001, and **** p<0.0001. (e) Fiber morphometry based on each fiber-type. Bar graph of aggregate fibers to the left and histogram to the right.

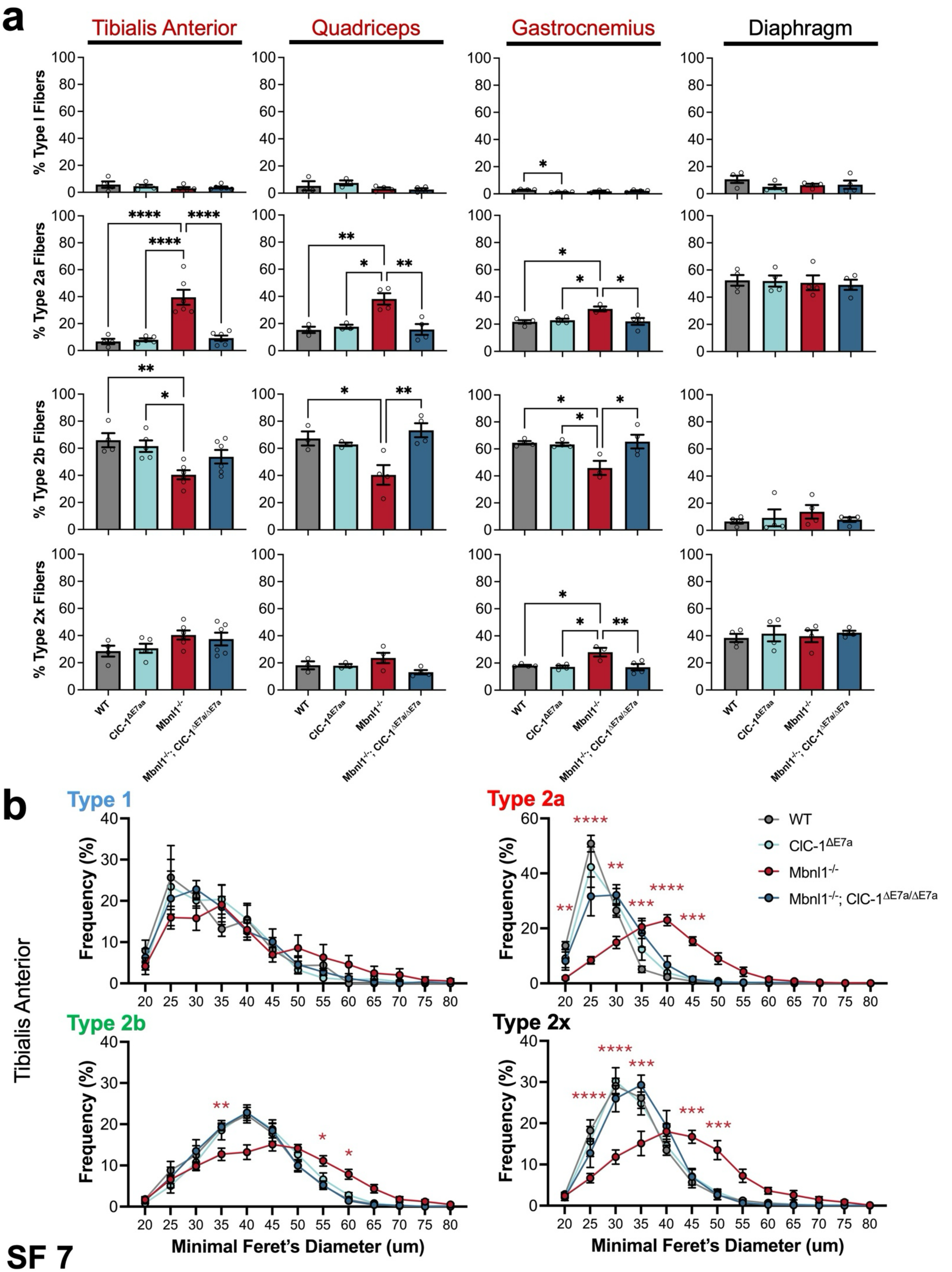

a
Tibialis Anterior
Quadriceps
Gastrocnemius
Diaphragm
% Type I Fibers
% Type 2a Fibers
% Type 2b Fibers
% Type 2x Fibers
WT
ClC-1ΔE7aa
Mbnl1-/-
Mbnl1-/-; ClC-1ΔE7a/ΔE7a
b
Type 1
Type 2a
Type 2b
Type 2x
Tibialis Anterior
Frequency (%)
Minimal Feret's Diameter (um)
WT
ClC-1ΔE7a
Mbnl1-/-
Mbnl1-/-; ClC-1ΔE7a/ΔE7a


SF 7

**Figure S7** Fiber type analysis across hindlimb and diaphragm muscles as well as TA fiber-type morphometry. (a) Fiber-type analysis by MyHC IHC for transverse slices from tibialis anterior (transposed from Fig. 5a), quadriceps, gastrocnemius, and diaphragm for WT, ClC-1$^{\Delta E7a/\Delta E7a}$, Mbnl1$^{-/-}$, Mbnl1$^{-/-}$ ; ClC-1$^{\Delta E7a/\Delta E7a}$ samples (4-6-months old). Data presented a mean ± SE with biological replicates shown as individual points. Comparisons with one-way ANOVA with post-hoc Tukey's multiple comparisons. (b) Histograms of minimum Feret's diameter for tibialis anterior fibers by fiber-type. Comparisons with two-way ANOVA (bin and genotype) with post-hoc multiple comparisons with stars representing intra-bin comparisons versus WT (red stars represent Mbnl1$^{-/-}$ versus WT). Multiplicity adjusted p-values shown as * $p < 0.05$, ** $p < 0.01$, *** $p < 0.001$, and $p < 0.0001$.

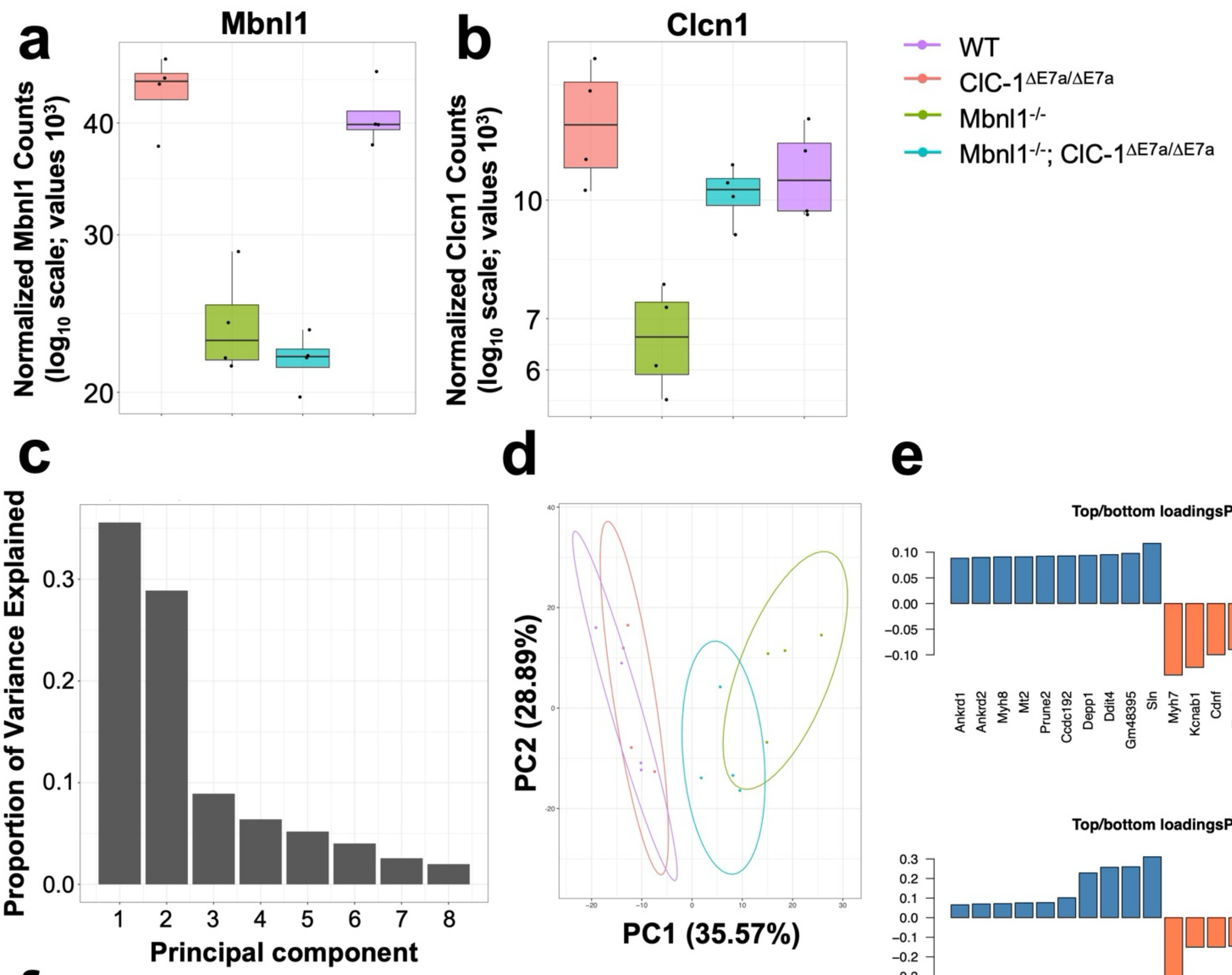


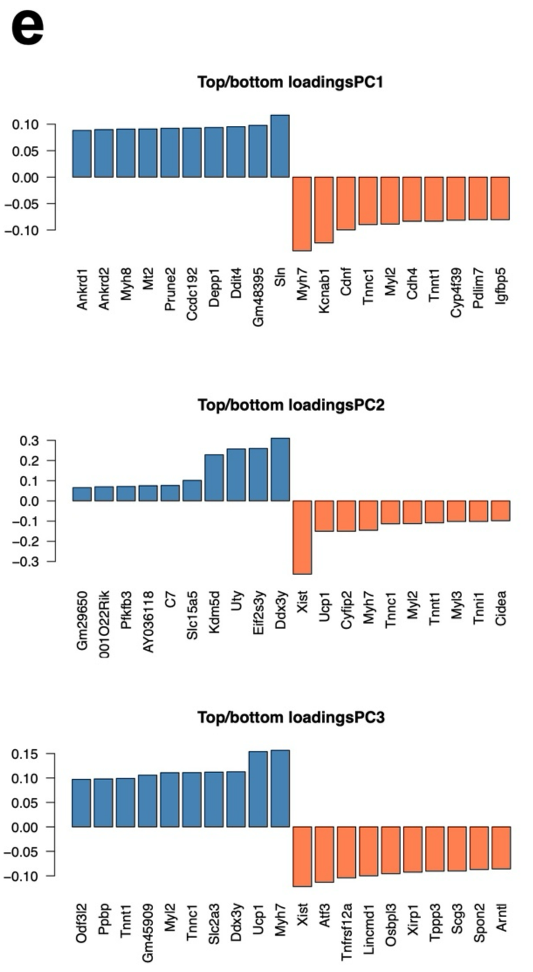


**f**

| Term | P-value | Adjusted p-value | Odds ratio | Combined score | Genes |
|---|---|---|---|---|---|
| Striated muscle contraction (GO:0006941) | 3.57E-09 | 3.14E-06 | 20.23514682 | 393.6007287 | Csrp3;Myh3;Tnnt1;Myl2;Myl3; Myh8;Tnni1;Myh6;Myh7 |
| Heart contraction (GO:0060047) | 1.52E-07 | 6.68E-05 | 21.08533984 | 331.0642343 | Csrp3;Actc1;Myl2;Myl3; Tnni1;Myh6;Myh7 |
| Myofibril assembly (GO:0030239) | 7.18E-07 | 1.45E-04 | 16.28497409 | 230.3788363 | Csrp3;Myh3;Tnnt1;Myl2;Ankrd1; Myh6;Lmod2 |
| Skeletal muscle contraction (GO:0003009) | 9.87E-07 | 1.45E-04 | 36.23809524 | 501.1275974 | Myh3;Tnnt1;Myh8;Tnni1;Myh7 |
| Cardiac muscle contraction (GO:0060048) | 9.88E-07 | 1.45E-04 | 21.83946981 | 301.988507 | Csrp3;Myl2;Myl3;Tnni1; Myh6;Myh7 |
| Sarcomere organization (GO:0045214) | 9.88E-07 | 1.45E-04 | 21.83946981 | 301.988507 | Csrp3;Myh3;Tnnt1;Ankrd1; Myh6;Lmod2 |

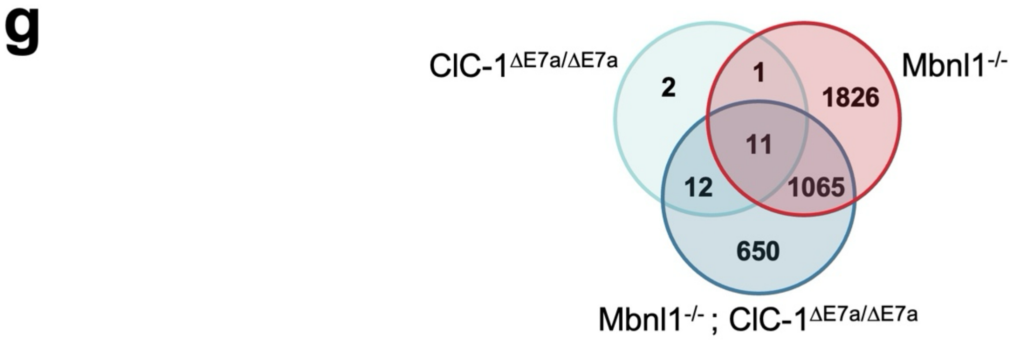


**SF 8**

**Figure S8** RNA-seq expression analysis. Normalized counts of reads for (a) Mbnl1 and (b) Clcn1 from RNAseq on approximately 4-6-month-old quadriceps tissue. Boxplots from pcaExplorer shown with individual values for each sample. (c) Scree plot of the variance accounted for by each principal component. (d) PCA plot with axes defined by PC1 and PC2. (e) Plot of top and bottom highest loading genes for PC1 through PC3. (f) Table displaying GO biological pathways analysis of top loading genes for PC1. (g) Venn diagram showing unique and shared dysregulated genes in ClC-1$^{\Delta E7a/\Delta E7a}$, Mbnl1$^{-/-}$, and Mbnl1$^{-/-}$ ; ClC-1$^{\Delta E7a/\Delta E7a}$ samples (compared to WT) with a threshold of absolute $\log_2$FC > 0.5 and adjusted p-value of < 0.05.

### MAJIQ voila output for several events across all samples

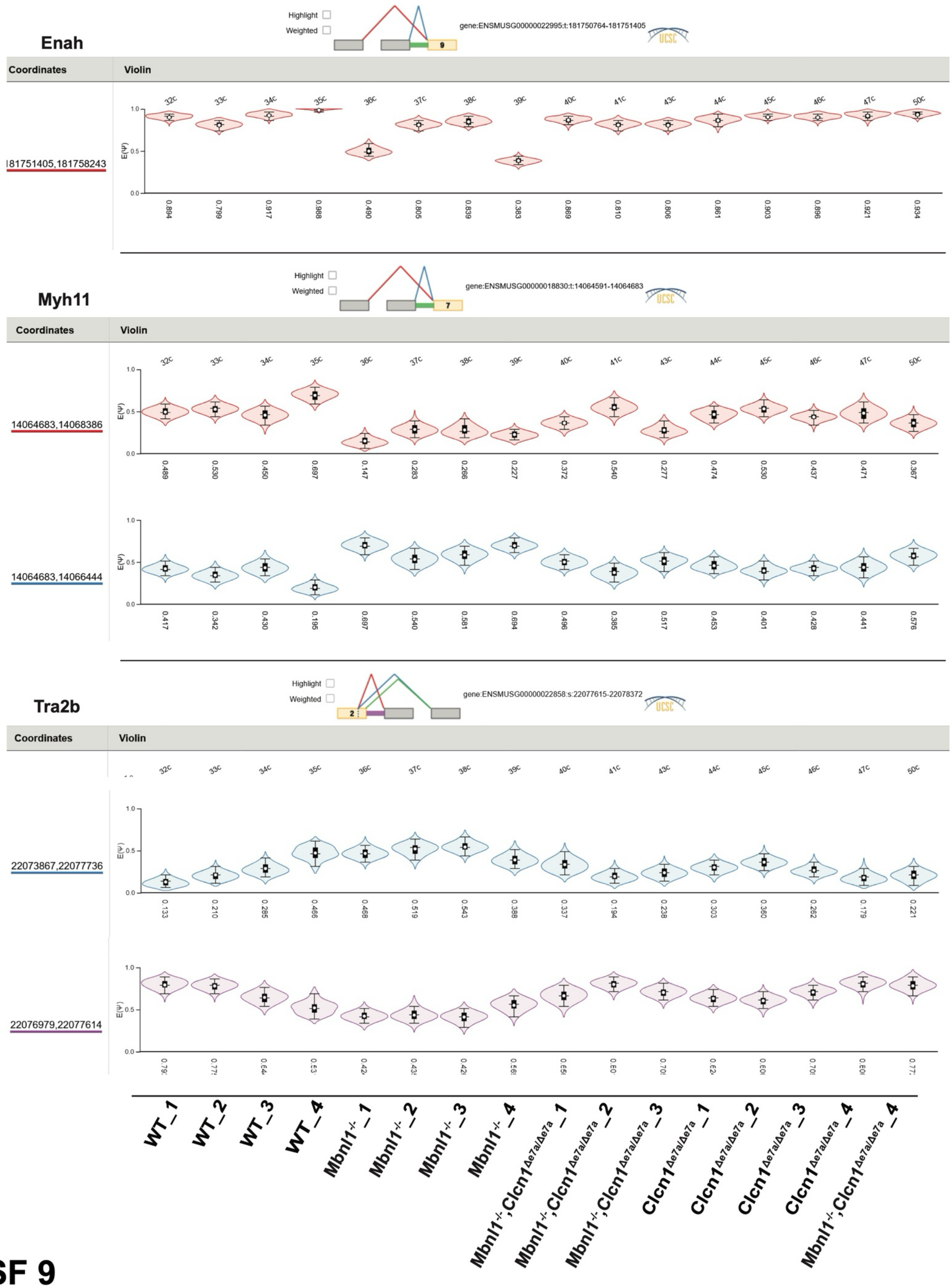


**SF 9**

**Figure S9** RNA-seq splicing analysis. Splicing analysis, MAJIQ output, for Enah, Myh11, and Tra2b.

**a**

| Gene | Use | F/R | Sequence (5' → 3') |
|---|---|---|---|
| Clcn1 | gDNA PCR<br>(E7a) | F<br>R | CCTAGCCATATCATCCCCTCTGTCCA<br>GCCCAGCAAATGAACAAATAGTGA |
| Clcn1[a] | RT-PCR<br>(E2→E3) | F<br>R | GGGAATGCCCAAGAAGATGGGCT<br>GCCATCAGGAGGCCCAGGAGCAC |
| Clcn1[b] | RT-PCR<br>(E5→E8) | F<br>R | TGAAGGAATACCTCACACTCAAGG<br>CACGGAACACAAAGGCACTG |
| Cacna1s[a] | RT-PCR<br>(E28→E30) | F<br>R | GAGATCCTTGGAATGTGTTTGACTTCCT<br>GGTTCAGCAGCTTGACCAGTCTCAT |
| Clcn1 | PacBio<br>(E1→E23) | F<br>R | 5AmMC6/GCAGTCGAACATGTAGCTGACTCAGGTCACTTGTACCA GCTACGGACTGC<br>5AmMC6/TGGATCACTTGTGCAAGCATCACATCGTAGCGTCCACTC TCTGACATGGG |
| Clcn1[c] | RT-qPCR<br>(ss-mRNA) | F<br>Probe<br>R | TGGAGACTGTGTGGGAGAC<br>5'6-FAM/CCAATTACT/ZEN/GCATAGCCCCCAGGT/3'IABkFQ<br>CATTTGGAAGGCTGGTAGGAG |
| Clcn1 | RT-qPCR<br>(pre-mRNA) | F<br>Probe<br>R | CTGTGTCATTCCCATTGTACCC<br>5'6-FAM/TGTCAAGAA/ZEN/AGACAGCTCCAACCCA/3'IABkFQ<br>TGGAGACTGTGTGGGAGAC |
| Tbp[d] | RT-qPCR<br>(reference) | F<br>Probe<br>R | CCAGAACTGAAAATCAACGCAG<br>5'TET/ACTTGACCT/ZEN/AAAGACCATTGCACTTCGT/3IABkFQ<br>TGTATCTACCGTGAATCTTGGC |

(a) Yadava *et al.* (2020), (b) Wheeler *et al.* (2007), (c) Mm.PT.58.8747621 (IDT), (d) Mm.PT.39a.22214839 (IDT)

**b**

**Primary Antibodies:**

| Anti- | Source | Product # | Isotype | Dilution |
|---|---|---|---|---|
| MyHC1 | DSHB | BA-D5 | MIgG2b | 1:40 |
| MyHC2a | DSHB | SC-71 | MIgG1 | 1:40 |
| MyHC2b | DSHB | BF-F3 | MIgM | 1:40 |
| Laminin | Sigma-Aldrich | L9393 | Rabbit poly | 1:1000 |

**Secondary Antibodies:**

| Anti- | Source | Product # | Fluorophore | Dilution |
|---|---|---|---|---|
| MIgG2b | Jackson Immuno. | 115-477-187 | DyLight 405 | 1:1000 |
| MIgG1 | Invitrogen | A-21124 | Alexa 568 | 1:1500 |
| MIgM | Invitrogen | A-21042 | Alexa 488 | 1:1500 |
| RIgG | Invitrogen | A-21245 | Alexa 647 | 1:1000 |

**ST 1**

**Table S1** Oligonucleotides and antibodies. (a) Sequences of primers and probes used for genotyping PCR reactions, RT-PCR splicing analysis, full-length *Clcn1* transcript amplification, and RT-qPCR. Primers were obtained from previous studies as well as IDT designer, as indicated. (b) Primary and secondary antibodies used for MyHC IHC.

| Genotype: | | | Linear Properties: | | | Instantaneous Current Density IV: | Steady-State Current Density IV: | Relative Open Probability: | | |
|---|---|---|---|---|---|---|---|---|---|---|
| | Age (days postnatal): | n | Capacitance (pF) | Rs (kOhms) | Tau (us) | Peak Instantaneous Current Density (pA/pF) | Peak Inward Current Density (pA/pF) | $P_{min}$ | k | $V_{1/2}$ (mV) |
| WT | 12 | 8 | 597±32 | 853±52 | 549±64 | -40.9±1.7 | -8.7±1.2 | 0.190±0.027 | 24.6±2.0 | -74.7±2.3 |
| ClC-$1^{\Delta E7a/\Delta E7a}$ | 12 | 11 | 494±25 | 1128±99 | 602±69 | -50.6±4.9 | -10.9±0.9 | 0.172±0.034 | 31.9±2.9 | -66.5±3.0 |
| Mbnl$1^{-/-}$ | 12 | 8 | 489±29 | 1059±51 | 560±40 | -8.5±0.8 | -1.3±0.1 | 0.179±0.018 | 19.9±1.6 | -59.9±1.7 |
| Mbnl$1^{-/-}$; ClC-$1^{\Delta E7a/\Delta E7a}$ | 12 | 8 | 502±19 | 1005±128 | 523±49 | -54.5±4.4 | -9.4±0.9 | 0.155±0.023 | 24.5±1.8 | -66.2±2.0 |
| WT | 18 | 14 | 1082±39 | 697±40 | 816.5±45 | -42.1±1.7 | -9.5±0.7 | 0.192±0.022 | 31.8±2.1 | -62.3±2.05 |
| ClC-$1^{\Delta E7a/\Delta E7a}$ | 18 | 17 | 985±31 | 657±29 | 703±36 | -63.5±4.2 | -18.4±1.8 | 0.171±0.041 | 39.6±3.9 | -64.9±3.74 |
| Mbnl$1^{-/-}$ | 18 | 16 | 831±30 | 703±31 | 631±37 | -25.3±1.2 | -5.3±0.4 | 0.133±0.038 | 31.2±2.7 | -75.2±3.19 |
| Mbnl$1^{-/-}$; ClC-$1^{\Delta E7a/\Delta E7a}$ | 18 | 14 | 851±42 | 742±36 | 677±35 | -56.4±2.7 | -13.8±1.1 | 0.139±0.030 | 33.8±2.6 | -72.2±2.58 |
| ADR | 18 | 7 | 1169±62 | 942±62 | 1129±81 | 0.031±0.06 | n/a | n/a | n/a | n/a |

**ST 2**

**Table S2** Whole-cell patch-clamp linear properties and fitting parameters. Table of linear properties and fitting properties for relative open probability for ClC-1 current recordings.
